# Compensatory variability in network parameters enhances memory performance in the *Drosophila* mushroom body

**DOI:** 10.1101/2021.02.03.429444

**Authors:** Nada Y. Abdelrahman, Eleni Vasilaki, Andrew C. Lin

## Abstract

Neural circuits use homeostatic compensation to achieve consistent behaviour despite variability in underlying intrinsic and network parameters. However, it remains unclear how compensation regulates variability across a population of the same type of neurons within an individual, and what computational benefits might result from such compensation. We address these questions in the *Drosophila* mushroom body, the fly’s olfactory memory center. In a computational model, we show that memory performance is degraded when the mushroom body’s principal neurons, Kenyon cells (KCs), vary realistically in key parameters governing their excitability, because the resulting inter-KC variability in average activity levels makes odor representations less separable. However, memory performance is rescued while maintaining realistic variability if parameters compensate for each other to equalize KC average activity. Such compensation can be achieved through both activity-dependent and activity-independent mechanisms. Finally, we show that correlations predicted by our model’s compensatory mechanisms appear in the *Drosophila* hemibrain connectome. These findings reveal compensatory variability in the mushroom body and describe its computational benefits for associative memory.

**Significance statement:** How does variability between neurons affect neural circuit function? How might neurons behave similarly despite having different underlying features? We addressed these questions in neurons called Kenyon cells, which store olfactory memories in flies. Kenyon cells differ among themselves in key features that affect how active they are, and in a model of the fly’s memory circuit, adding this inter-neuronal variability made the model fly worse at learning the values of multiple odors. However, memory performance was rescued if compensation between the variable underlying features allowed Kenyon cells to be equally active on average, and we found the hypothesized compensatory variability in real Kenyon cells’ anatomy. This work reveals the existence and computational benefits of compensatory variability in neural networks.

## Introduction

Noise and variability are inevitable features of biological systems. Neural circuits achieve consistent activity patterns despite this variability using homeostatic plasticity: because neural activity is governed by multiple intrinsic and network parameters, variability in one parameter can compensate for variability in another to achieve the same circuit behaviour [1–5]. This phenomenon of compensatory variability has typically been addressed from the perspective of consistency of neural activity across individual animals [6, 7] or over an animal’s lifetime, in the face of circuit perturbations [8–11]. However, less attention has been paid to potential benefits of maintaining consistent neuronal properties across a population of neurons within an individual circuit.

Indeed, previous work has emphasized the benefits of neuronal heterogeneity rather than neuronal homogeneity [12–14]. Of course, different neuronal classes encode different information (e.g., visual vs. auditory neurons, or ON vs. OFF cells). Yet even in populations that ostensibly encode the same kind of stimulus, like olfactory mitral cells, heterogeneity of neuronal excitability can increase the information content of their population activity [15–17]. In addition, heterogeneity in neuronal time scales can improve learning in neural networks [18, 19]. In what contexts and in what senses might the opposite be true, i.e., when does neuronal similarity provide computational benefits over neuronal diversity? And what mechanisms could enforce neuronal similarity in the face of inter-neuronal variability?

Here we address these questions using olfactory associative memory in the mushroom body of the fruit fly *Drosophila*. Flies learn to associate specific odors with salient events (e.g., food or danger). These olfactory associative memories are stored in the principal neurons of the mushroom body, called Kenyon cells (KCs), as modifications in KCs’ output synapses [20–22] (reviewed in [23]). Because learning occurs at the single output layer, the nature of the odor representation in the KC population is crucial to the fly’s ability to learn to form distinct associative memories for different odors. In particular, the fact that KCs respond sparsely to incoming odors (≈ 10% per odor) [24] allows different odors to activate unique, non-overlapping subsets of KCs and thereby enhances flies’ learned discrimination of similar odors [25].

A potential problem for this sparse coding arises from variability between KCs. KCs receive inputs from second-order olfactory neurons called projection neurons (PNs), with an average of ≈ 6 PN inputs per KC, and typically require simultaneous activation of multiple input channels in order to spike [26], thanks to high spiking thresholds and feedback inhibition [25, 27]. However, there is substantial variation across KCs in the key parameters controlling their activity, such as the number of PN inputs per KC [28], the strength of PN-KC synapses, and KC spiking thresholds [27]. Intuitively, such variation could lead to a situation where some KCs with low spiking thresholds and many or strong excitatory inputs fire indiscriminately to many different odors, while other KCs with high spiking thresholds and few or weak excitatory inputs never fire; KCs at both extremes are effectively useless for learning to classify odors, even if overall only 10% of KCs respond to each odor. However, it remains unclear whether biologically realistic inter-KC variability would affect the mushroom body’s memory performance, and what potential strategies might counter the effects of inter-KC variability.

Here we show in a rate-coding model of the mushroom body that introducing experimentally-derived inter-KC variability into the model substantially impairs its memory performance. This impairment arises from decreased dimensionality of the KC population activity and increased similarity between KC responses to different odors, ultimately arising from the variability in average activity among KCs. However, memory performance can be rescued by compensating away variability in KC activity while preserving the experimentally observed variation in the underlying parameters. This can occur through activity-dependent homeostatic plasticity or direct correlations between key parameters like number vs. strength of inputs. Finally, we analyze the hemibrain connectome to show that indeed, the number of PN inputs per KC is inversely correlated with the strength of each input, while the strength of inhibitory inputs is correlated with the total strength of excitatory inputs. Thus, we show both the existence and computational benefit of compensatory variability in mushroom body network parameters.

## Results

### Realistic inter-KC variability impairs memory performance

To study how variability between KCs might affect the fly’s olfactory memory performance, we modelled the mushroom body as a rate-coding neural network (Fig. 1). To simulate the input activity from PNs, we modeled their activity as a saturating non-linear function of activity of the first-order olfactory receptor neurons (ORNs) (see Methods; [29]). We applied this function to the recorded odor responses of 24 different olfactory receptors [30] to yield simulated PN activity, as has been done in many computational studies of fly olfaction [31–34]. To simulate variability in PN activity across different encounters with the same odor, we created several ‘trials’ of each odor and added Gaussian noise to PN activity, following the coefficients of variation reported in [35]. To increase the number of stimuli beyond the 110 recorded odors in [30], we generated odor responses in which the activity of each PN was randomly sampled from that PN’s activity across the 110 odors used in [30] (results were similar with the ‘real’ 110 odors; see Methods and below).

**Figure 1:**
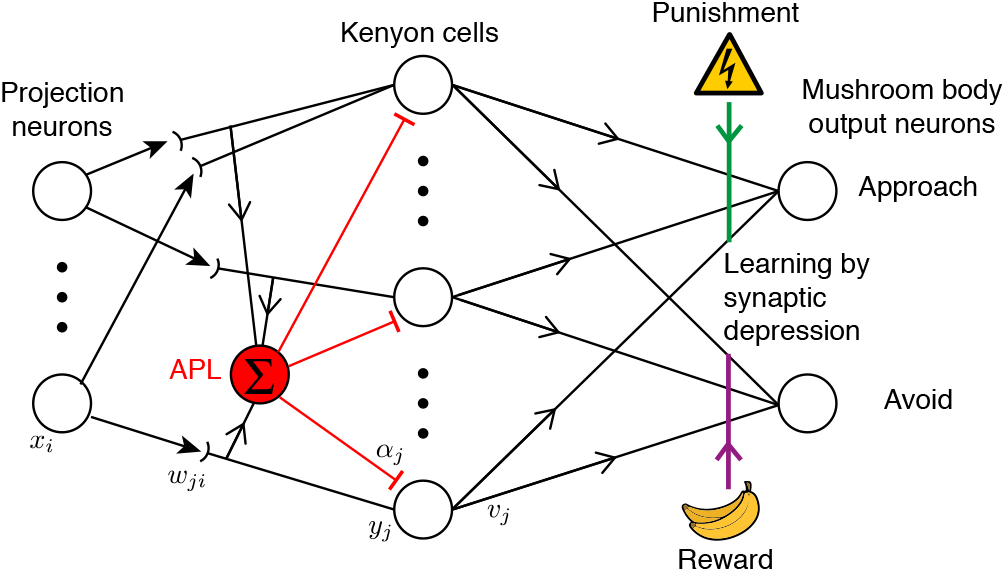
Schematic for the mushroom body network model. Projection neurons in the input layer relay the odor responses, *x_i_*, downstream to the Kenyon cells (*y_j_*). Kenyon cells connect randomly to the projection neurons with synaptic weights *w_ji_* and receive global inhibition from the APL neuron with weight *α_j_*. Learning occurs when dopaminergic neurons (DANs) carrying punishment (reward) signals from the environment depress the synapses (*v_j_*) between the active Kenyon cells and the mushroom body output neurons (MBONs) that lead to approach (avoidance) behavior.

Each KC in our model received excitatory input from a randomly selected set of *N* PNs, each with strength *w*. A KC’s response to each odor was the sum of excitatory inputs minus inhibition, minus a spiking threshold *θ*; if net excitation was below the threshold, the activity was set to zero. Inhibition came from the feedback interneuron APL (‘Anterior Paired Lateral’), which is excited by and inhibits all KCs [25]. To avoid simulating the network in time, we simplified the feedback inhibition into pseudo-feedforward inhibition, in which APL’s activity was the sum of all post-synaptic excitation of all KCs (without the KCs’ threshold applied); we based this simplification on the fact that KCs and APL form reciprocal synapses with each other on KC dendrites (i.e., before the KCs’ spike initiation zone), and APL activity is somewhat spatially restricted between KC axons and dendrites [36].

Learning in flies occurs when KCs (responding to odor) are active at the same time as dopaminergic neurons (DANs, responding to ‘reward’ or ‘punishment’); the coincident activity modifies the output synapse from KCs onto mushroom body output neurons (MBONs) that lead to behavior (e.g., approaching or avoiding an odor). Typically, the output to the ‘wrong’ behavior is depressed: for example, pairing an odor with electric shock weakens the output synapses from KCs activated by that odor onto MBONs that lead to ‘approach’ behavior [21, 22, 37, 38] (reviewed in [23]). We simulated this plasticity using a simplified architecture with only two MBONs, one for ‘approach’ and one for ‘avoid’. The input odors were randomly divided: half were paired with punishment and half with reward. During training, KCs activated by rewarded odors weakened their synapses onto the ‘avoid’ MBON, while KCs activated by punished odors weakened their synapses onto the ‘approach’ MBON (depression by exponential decay; see Methods). The fly’s behavior then depended probabilistically (via a softmax function; see Eq. 21, Methods) on whether the ‘avoid’ or ‘approach’ MBON’s was greater, and the model’s accuracy in learning was scored as the fraction of correct decisions for unseen noisy variants of the trained odors (i.e., avoiding punished odors and approaching rewarded odors).

To test the effect of realistic inter-KC variability on this model, we introduced variability step-by-step. We first tested the performance of the model holding constant across all KCs the 3 parameters *N* (number of PN inputs per KC), *w* (strength of each PN-KC connection) and *θ* (KC spiking threshold). Then we added inter-KC variability step-by-step: first varying only one out of 3 parameters, then 2 out of 3, then all 3 parameters (thus 8 possible models). Inter-KC variability in *N*, *w* and *θ* followed experimentally measured distributions (Fig. 2A1-3) [27, 28]. Increasing inter-KC variability systematically degraded the model’s performance when tested on 100 input odors: the more variable parameters there were, the worse the performance (Fig. 2B). In the two extreme cases, the model with all 3 parameters fixed performed at 72.5% accuracy while the model with all 3 parameters variable performed at 64% accuracy. This performance trend was the same when these 8 models were trained and tested on the real input odors responses from [30] (78.1% v. 63.9% Fig. S1).

**Figure 2:**
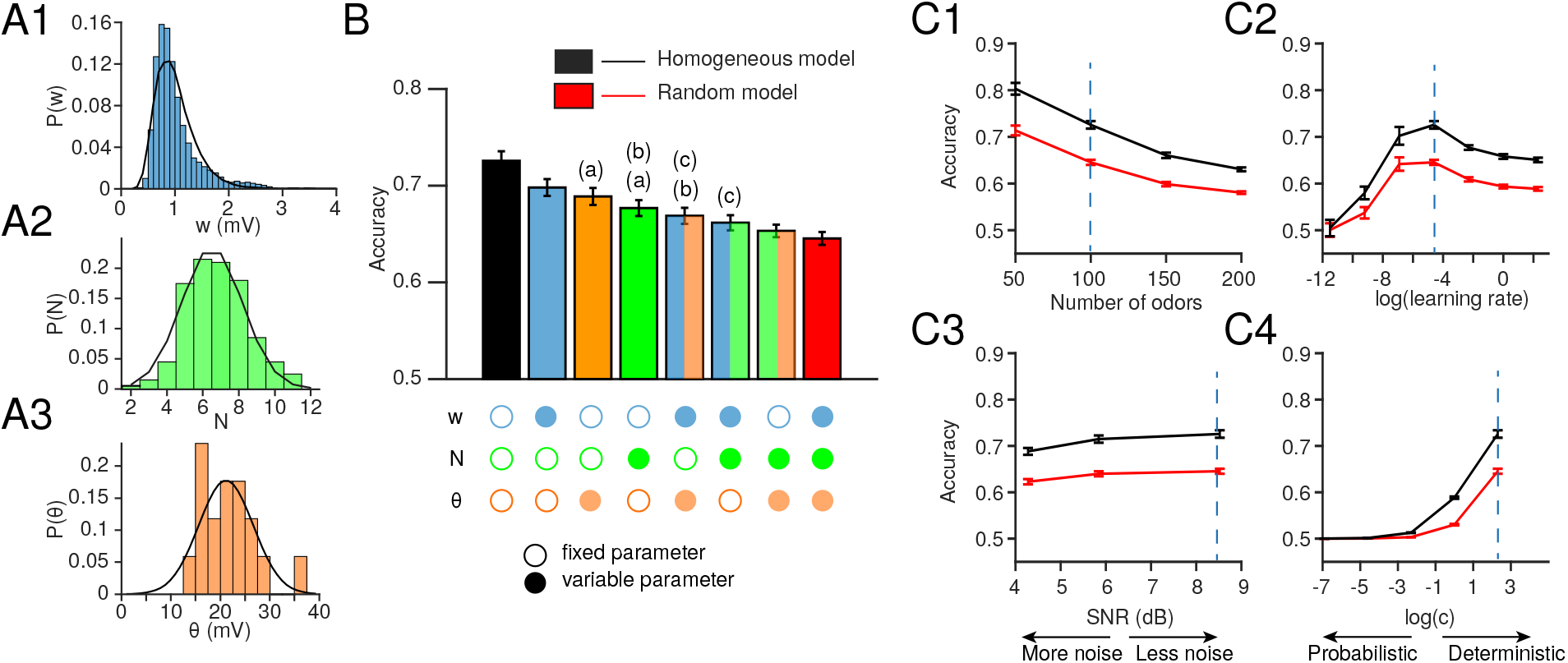
Inter-KC variability in *w*, *N* and *θ* degrades the model fly’s memory performance. (**A**) Histograms of the experimentally measured distributions for: (**A1**) *w* (amplitude of spontaneous excitatory postsynaptic potentials in KCs, mV; data from [27]), (**A2**) *N* (number of PN inputs per KC, measured as the number of dendritic ‘claws’; data from [28]), (**A3**) *θ* (spiking threshold minus resting potential, mV; data from [27]). The overlaid black curves show log-normal (*w*) and Gaussian (*N*, *θ*) fits to the data. (**B**) The model fly’s memory performance (given 100 input odors), varying the parameters step by step. Fixed and variable parameters are shown by empty and filled circles, respectively. The homogeneous model (all parameters fixed; black) performs the best and the random model (all parameters variable; red) performs the worst. All bars are significantly different from each other unless the share the same letter annotations (a, b, etc.), *p* < 0.05 by Wilcoxon signed-rank test (for matched models with the same PN-KC connectivity) or Mann-Whitney test (for unmatched models with different PN-KC connectivity, i.e., fixed vs. variable *N*), with Holm-Bonferroni correction for multiple comparisons (full statistics in Table S1). *n* = 25 model instances with different random PN-KC connectivity, error bars show twice the standard error of the mean (95.4% confidence interval). (**C**) The performance trend is consistent over a range of different conditions: (**C1**) number of input odors, (**C2**) the learning rate used to learn the optimum weights between KCs and MBONs, (**C3**) amount of noise in PN activity (measured by signal-to-noise ratio, SNR), (**C4**) the indeterminacy in the decision making, quantified by log(c), where c is the constant in the soft-max function (Eq. 21). The vertical dotted lines indicate the conditions used in panel B.

To test whether this effect is robust to different learning and testing conditions, we tested the two extreme cases while varying the numbers of input odors to be classified, the amount of noise in PN activity, the learning rate at the KC-MBON synapse (the two models might have different optimal learning rates: *η* in Eq. 20), or the indeterminacy of the fly’s decision making (*c* in the softmax equation, Eq. 21). In every case, the model with all parameters fixed (which we call the ‘homogeneous’ model) consistently outperformed the model with all parameters variable (which we call the ‘random’ model) (Fig. 2C1-4). These results indicate that biologically realistic variability in KC network parameters impairs the network’s ability to classify odors as rewarded vs. punished.

### Realistic inter-KC variability reduces separation between KC odors representations

We next asked what features of the KC population odor representation might account for the worse performance of the random model compared to the homogeneous model. Learning the optimal KC-MBON weights to correctly classify the rewarded versus punished odors is equivalent to finding a hyper-plane (in 2000-dimensional space) to separate KC responses to rewarded odors from those to punished odors. Therefore, a model with better separability between KC odor representations would find a better separating hyper-plane, and have better performance in classifying unseen noisy variants of the trained odors. We measured separability using a variety of metrics.

We first asked whether odors are more widely separated in KC coding space in the homogeneous model, using angular distance, a scale-insensitive distance metric (see Methods). For each odor, we took the centroid of KC responses to the noisy variants of that odor, and for each pair of odors, we measured the angular distance between their respective centroids (Fig. 3A1). Indeed, the angular distance between odors (averaged across all odor pairs) was larger in the homogeneous model (Fig. 3A2), which matched the higher accuracy (Fig. 3A3, where each dot represents one instantiation of the network). This difference also extended to the angular distance between the centroids of the groups of odors randomly assigned to be rewarded and punished (Fig. 3B2), suggesting that the greater inter-odor distances in the homogeneous model make it easier to draw a hyper-plane separating the rewarded and punished odors.

**Figure 3:**
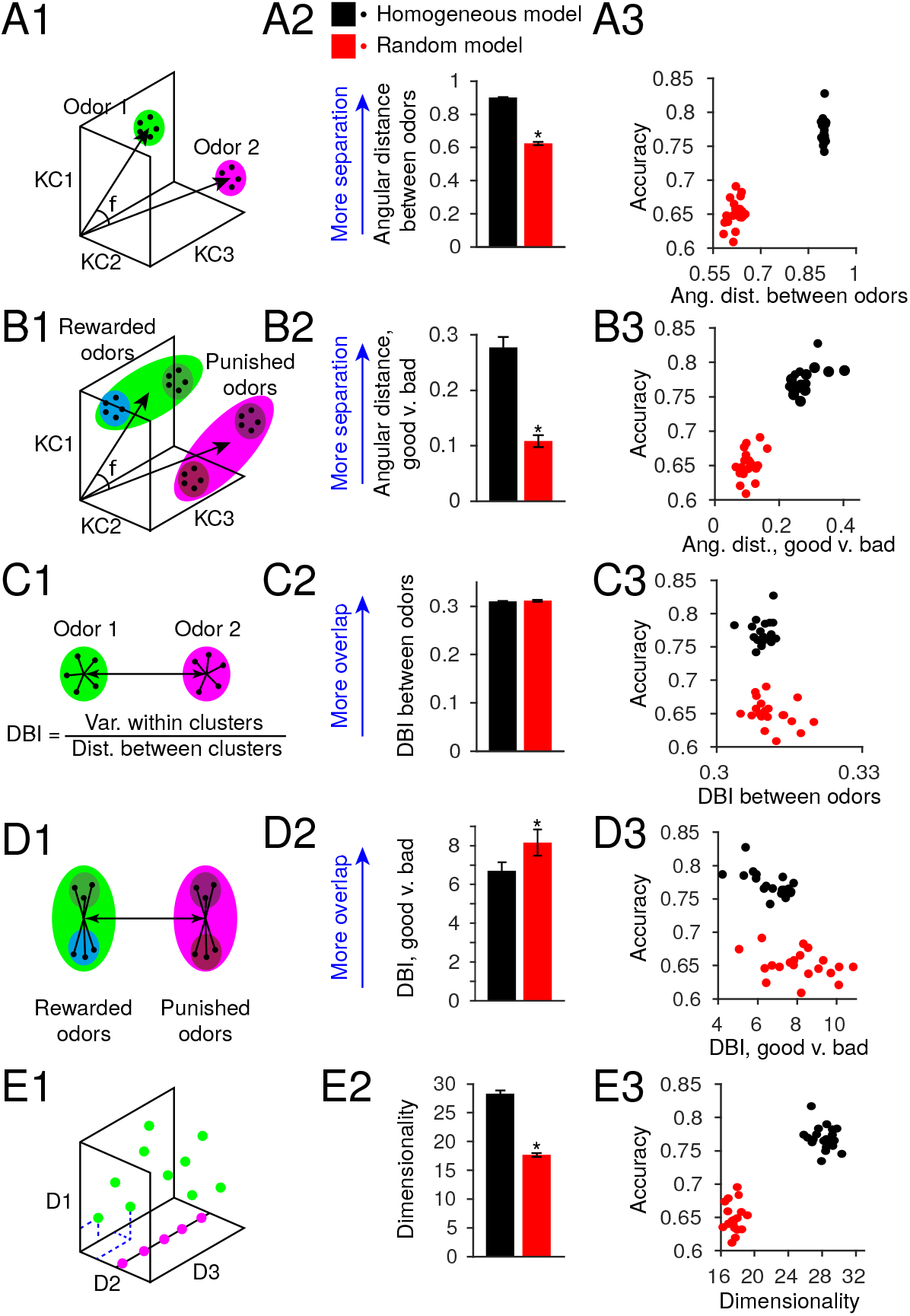
Inter-KC variability in *w*, *N* and *θ* reduces separability of KC odor representations. **Left column**: **(A1-D1)** show schematic illustrations of separability metrics: angular distance between individual odors (**A1**) or between rewarded vs. punished odors (**B1**), and DBI between individual odors (**C1**) or between rewarded vs. punished odors (**D1**). **(E1)** shows the dimensionality of a system with 3 variables. The system with its states scattered throughout 3D space (green) has dimensionality 3 while the system with all states on a single line (magenta) has dimensionality 1. **Middle column**: Separability metrics for the homogeneous (black) and random (red) models. Compared to the random model, the homogeneous model has higher angular distance **(A2,B2)**, similar DBI between odors **(D2)** but lower DBI between rewarded vs. punished odors **(D2)**, and higher dimensionality **(E2)**. Error bars show twice the standard error of the mean. * difference between homogeneous and random models, *p* < 0.05, Mann-Whitney test (full statistics in Table S1). **Right column**: **(A3-E3)** Scatter plots show performance vs. separability metric in the respective rows, calculated in *n*=25 random instantiations of the network.

However, the separability of clusters of noisy variants of odors might depend not only on the distance between their centroids, but also on their variability. For instance, two clusters of noisy variants with well separated centroids might overlap if the data points in the clusters are not tightly packed. Therefore, we next measured the quality of clustering in each model using the Davies-Bouldin Index (DBI). DBI measures the variance within clusters divided by the distance between the centroids of each cluster [39], so high DBI means more overlapping, less separable clusters. When we calculated DBI using different pairs of odors (Fig. 3C1), treating each odor (with its noisy variants) as its own cluster, DBI values were similar in the random and homogeneous models (Fig. 3C2), suggesting that poor performance in the random model was not explained by poor clustering of noisy variants (Fig. 3C3). (The DBI was slightly higher in the random model using the original odors from [30]: Fig. S1). However, DBI was higher in the random model when considering the two clusters of all rewarded odors vs. all punished odors (Fig. 3D1-2), and showed a weak inverse correlation with memory performance (Fig. 3D3) (note that each instantiation of the network received the same odors but different random reward/punishment assignments). These results suggest that in the homogeneous model (compared to the random model), odor representations are arranged in KC coding space in a way to allow punished and rewarded odors to be more easily separated.

We hypothesized that odor responses in the homogeneous model are more separable because they are arranged across more dimensions in KC coding space, allowing them more degrees of freedom. We quantified dimensionality according to [40]. Dimensionality of a dynamic system is the number of independent dimensions that define the system’s response to a given input. For example, if a system nominally has 3 dimensions but all its responses lie on a straight line, its dimensionality is only 1, in contrast to a system whose responses are distributed throughout the 3-dimensional space (Fig. 3E1). We found that KC responses in the homogeneous model had a significantly higher dimensionality than those in the random model (Fig. 3E2), matching the higher performance in the homogeneous model (Fig. 3E3). Together, these metrics indicate that introducing the realistic inter-KC variability in *w*, *N*, and *θ* worsens the performance of the network by reducing the dimensionality (and thus separability) of KC odor representations.

### Realistic inter-KC variability weakens specialization of KC responsiveness

We hypothesized that the lower dimensionality of the random model might arise because fewer KCs provide useful odor identity information when some are indiscriminately active while others are completely silent. Sparse coding requires sparseness in two dimensions: population sparseness (each stimulus activates few neurons) and lifetime sparseness (each neuron responds to few stimuli) [41]. While our models enforced population sparseness by scaling inhibition and spiking thresholds to achieve a coding level (fraction of cells active per odor) of 0.1 (averaged across all odors), they did not enforce any particular lifetime sparseness. In an extreme case, a model could have very consistent population sparseness with a coding level of 0.1 for all odors, simply by having the same 10% of cells responding to every odor and the other 90% being completely silent. In this case, none of the cells would provide any useful information about odor identity and dimensionality would be 0. We asked whether a less extreme version of this problem could explain the lower dimensionality and memory performance of the random model.

We measured the lifetime sparseness of KCs in the homogeneous and random models. Lifetime sparseness quantifies how specialized a cell is to particular input stimuli: 1 means a cell fires to one stimulus and no other stimuli, while 0 means it fires equally to all stimuli. A cell that fires to no stimuli has an undefined sparseness (see Methods). The homogeneous model had fairly consistent lifetime sparseness values, with almost 90% of KCs having a lifetime sparseness between ~0.85 and 1. In contrast, the random model had KCs with much more variable lifetime sparseness, with a long tail of KCs with low sparseness (below 0.7) and more than 40% of KCs having undefined sparseness (i.e., completely silent). The contrasting distributions of lifetime sparseness can seen in the cumulative distribution functions (cdfs) of lifetime sparseness in Fig. 4A, in how the steep curve of the homogeneous model and the shallow curve of the random model cross each other. This result can also be seen in the larger standard deviation of lifetime sparseness across KCs in the random model (Fig. 4B). The silent KCs can be seen as the fraction of missing KCs needed for the cdf curves to reach 1; the random model has many more silent KCs than the homogeneous model (Fig. 4A). Because silent KCs are useless for odor identity coding, a high number of silent KCs corresponds to low dimensionality of KC odor representations (Fig. 4C).

**Figure 4:**
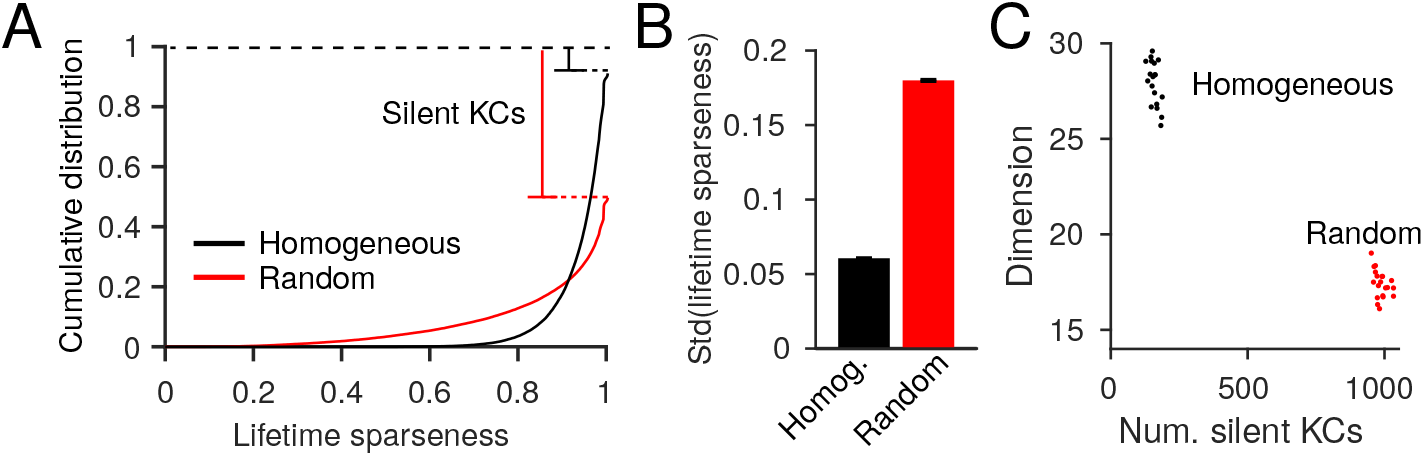
Inter-KC variability increases variability of lifetime sparseness and fraction of silent KCs. **(A)** Cumulative distribution function (cdf) of the lifetime sparseness of KCs in the homogeneous (black) and random (red) models, across 20 instantiations of the network. The gap between 1.0 and the top of the cdf represents silent KCs (lifetime sparseness undefined). **(B)** The random model has larger standard deviation in lifetime sparseness among KCs. Error bars show twice the SEM, *n* = 20 random instantiations of the network. Bars are different, *p* < 0.05, Mann-Whitney test (see Table S1). **(C)** Number of silent KCs plotted versus the dimensionality of KCs; each dot is one random model instance.

### Compensatory tuning of KC parameters rescues memory performance

Because the central problem for memory performance in the random model was inter-KC variability in average levels of activity, we hypothesized that performance could be rescued in models where KCs could achieve roughly equal activity across the population, while still preserving experimentally realistic variability in spiking thresholds and number/strength of excitatory inputs.

#### Parametric tuning of excitatory input weights

First, we tested a model that equalizes KC activity indirectly, by making parameters compensate for each other in an activity-independent way. In particular, we modeled KCs as adjusting input synaptic weights (*w*) to compensate for variability in spiking threshold (*θ*) and number of PN inputs (*N*). Thus, an individual KC with low *θ* or high *N* would have low *w*, while a KC with high *θ* or low *N* would have high *w*. We simulated these correlations 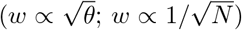 constrained by experimental data. To do this, we sampled *N* and *θ* from the distributions in Fig. 2A, and sampled *w* from a posterior compensatory distribution, *P* (*w* | *n, θ*), whose overall shape across all KCs was constrained to be the same as the experimental *P* (*w*) in Fig. 2A1 but which was composed of multiple distributions of *P* (*w*) for different values of *N* and *θ*. For example, a KC with a relatively high *n* = 7 would sample its weights from a *P* (*w*) shifted to the left (lower *w*) (Fig. 5A1, dashed lines), while a KC with a relatively high *n* = 2 would sample its weights from a *P* (*w*) shifted to the right (higher *w*) (Fig. 5A1, solid lines). The same would be true for different values of *θ* (Fig. 5A1, different shadings). We fitted these component *P* (*w*) curves so that with experimentally observed distributions of *N* and *θ*, the sum of the components would produce the experimentally observed distribution of *w* across all KCs (see Methods). (Note that this algorithm is not meant to describe an actual biological mechanism, merely to create correlations between *w* vs. *N* and *θ* while constraining the parameters to experimentally realistic distributions. Biologically, such correlations could arise through several mechanisms; see Discussion.) This compensatory mechanism rescued the fly’s performance, producing significantly higher accuracy at classifying odors than the random model (Fig. 5B, cyan bars), likely resulting from the higher dimensionality of KC representations (Fig. 5C) and reduced variability in KC lifetime sparseness (Fig. 5D).

**Figure 5:**
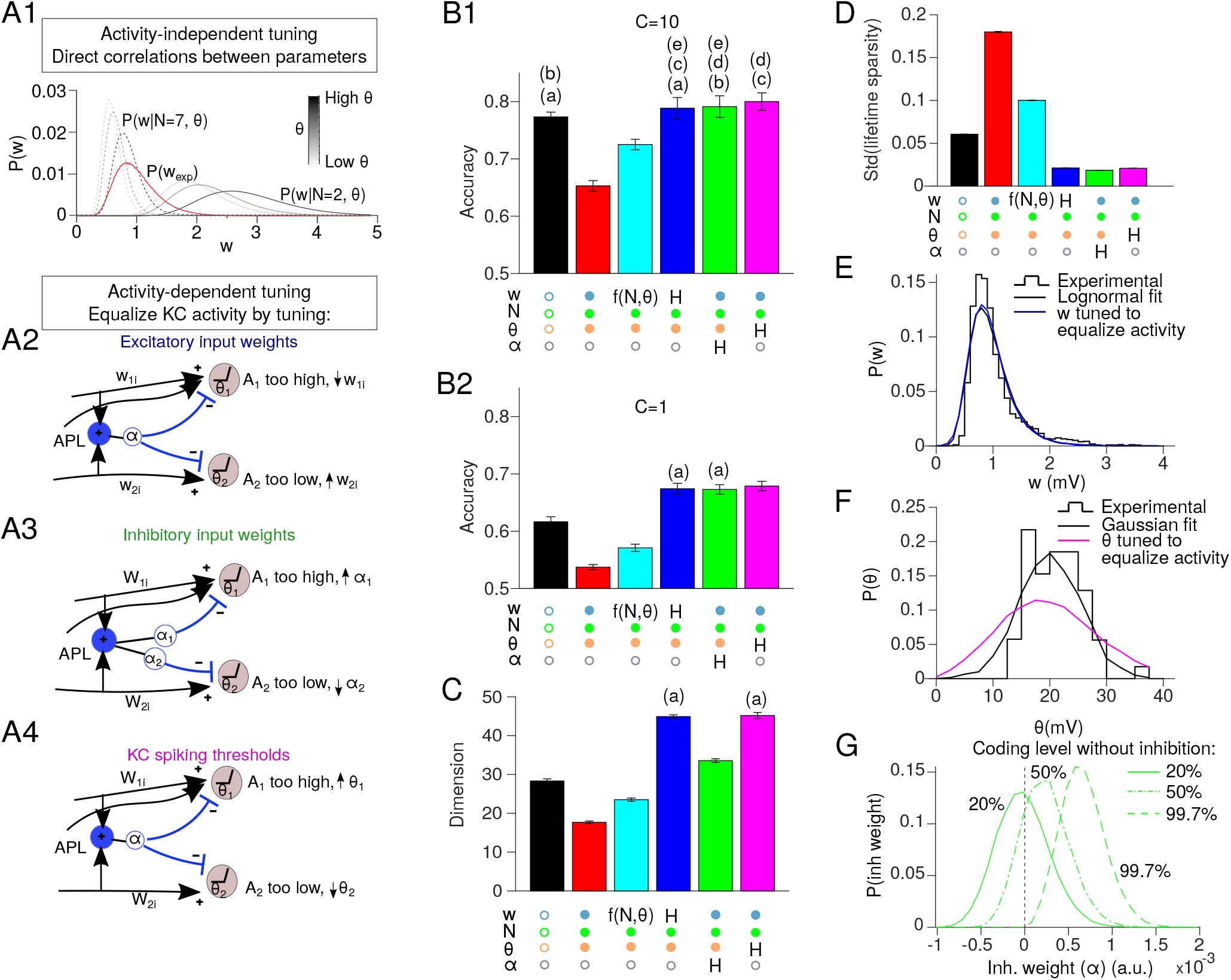
Compensation in network parameters rescues memory performance. **(A)** Schematics of different compensation methods. **(A1)** Lognormal fit of experimental distribution of the synaptic weights *P* (*w_exp_*) (red), and its component distributions for different *w* and *θ*, *P* (*w* | *N, θ*), for high *N* = 7 (dotted) or low *N* = 2 (solid). Shadings of gray indicate different values of *θ*. **(A2-4)** Mechanisms for activity-dependent homeostatic compensation. Overly active KCs weaken excitatory input weights (*w_ji_*, A2), strengthen inhibitory input weights (*α_j_*, A3), or raise spiking thresholds (*θ_j_*, A4). Inactive KCs do the reverse. **(B1)** Compensation rescues performance, alleviating the defect caused by inter-KC variability in the random model (red) compared to the homogeneous model (black), whether compensation occurs by setting *w* according to *N* and *θ* (cyan; A1), using activity-dependent homeostatic compensation to adjust excitatory weights (dark blue; A2), inhibitory weights (green; A3) or spiking thresholds (magenta; A4). **(B2)** Differences between models are more apparent when the task is more difficult due to more stochastic decision-making (*c* = 1 instead of *c* = 10 in the softmax function in Eq. 21). **(C-D)** Dimensionality of KC representations (C) and standard deviation of KC lifetime sparseness (D) in the models described above. Activity-dependent models have the highest dimensionality and lowest variability in KC sparseness. *n* = 20 model instances with different random PN-KC connectivity. Error bars show two times the SEM, i.e., 95.4% confidence interval. All bars are significantly different from each other unless the share the same letter annotations, *p* < 0.05, by Wilcoxon signed-rank test (for matched models with the same PN-KC connectivity) or Mann-Whitney test (for unmatched models with different PN-KC connectivity, i.e., fixed vs. variable *N*), with Holm-Bonferroni correction for multiple comparisons (full statistics in Table S1). Annotations below bars indicate whether parameters were fixed (empty circle), variable (filled circle), or variable following a compensation rule (‘H’ for homeostatic tuning, *f* (*N, θ*) for parametric tuning). (**E**) KC excitatory input synaptic weights (*w*) after tuning to equalize average activity (blue) follow a similar distribution to experimental data (black, from Fig. 2A1) (**F**) KC spiking thresholds (*θ*) after tuning to equalize average activity (magenta) have wider variability than the experimental distribution (black, from Fig. 2A3). (**G**) Tuning KC inhibitory weights (*α*) to equalize average activity requires many inhibitory weights to be negative, unless the coding level without inhibition is as high as 99.7%.

#### Activity-dependent tuning of KC parameters

We next tested compensatory mechanisms based on activity rather than explicit correlations between network parameters. Here, each KC has the same desired average activity level across all odors, *A*_0_ (with a tolerance of ±6%). We tested three models, each of which equalized average KC activity *A*_0_ by tuning a different parameter: input excitatory weights (*w*), inhibitory weights (*α*), or spiking thresholds (*θ*). The non-tuned parameters followed the distributions in Fig. 2A (inhibitory weights were constant when non-tuned), while individual KCs adjusted the tuned parameter according to whether their activity was too high or too low. For example, a relatively highly active KC (whether because it has high *w* or *N*, low *θ*, or simply receives input from highly active PNs) would scale down its excitatory weights (Fig. 5A2), scale up its inhibitory weights (Fig. 5A3), or scale up its spiking threshold (Fig. 5A4). Likewise, a relatively inactive (or indeed totally silent) KC would do the reverse (see Methods for details of the update rules underlying the homeostatic tuning and discussion of variant update rules shown in Figs. S3,S4).

All three homeostatic models performed as well as the homogeneous model (Fig. 5B1, blue, green, magenta bars), and indeed even out-performed the homogeneous model when decision-making was more stochastic (lower value of *c* in the softmax function; Fig. 5B2). The more stochastic decision-making makes the task more difficult and thus brings out the enhanced coding by the homeostatic models. Indeed, the dimensionality of KC odor representations in the homeostatic models was even higher than that in the homogeneous model (Fig. 5C), and the variability in KC lifetime sparseness was even lower (Fig. 5D).

What distributions of excitatory weights, inhibitory weights, or spiking thresholds emerge after activity-dependent tuning to equalize KC activity? Do they match experimentally observed distributions? Tuning excitatory weights led to a distribution fairly similar to the approximately log-normal experimentally observed distribution of EPSP amplitudes (Fig. 5E). Tuning spiking thresholds led to a distribution with greater variance than the experimental distribution, although with a qualitatively similar Gaussian shape (Fig. 5F). This larger variance of thresholds suggests that natural variation of *θ* is too small, on its own, to equalize KC activity given the variation in the number/strength of excitatory inputs.

The tuned distribution of inhibitory weights differed even more strongly from experimental results. While there are no experimental measurements of inhibitory weights, equalizing KC activity by tuning inhibitory weights required many of them to be negative (Fig. 5G), which is unrealistic, because negative inhibition is actually excitation, and there are no reports of GABAergic excitation of KCs [42].

Why did our model require negative inhibition? This result can be understood by considering one of the model’s constraints: that inhibition is only strong enough to reduce the fraction of active KCs by half, i.e., 10% of KCs are active on average in normal flies, while 20% of KCs are active if inhibition is blocked (based on results from [25]). Because the average threshold must be high enough that 80% of KCs are silent on average even without inhibition, the wide variation in thresholds and excitation means that many KCs will have excitation so weak, and thresholds so high, that no stimulus could ever drive them above threshold, even in the absence of inhibition. For inhibition to compensate for inactivity even in the absence of inhibition, it must become negative (i.e., excitatory) in these weakly-activated KCs. In contrast, the models that tune excitatory weights or thresholds do not face this problem, because inactive KCs can simply increase their excitatory weights or decrease their thresholds. The central problem for the inhibitory plasticity model is that inhibition is not a strong enough force in our system to balance out variable excitation and thresholds on its own without becoming negative. Indeed, if we relax the constraint that coding level be 0.2 without inhibition, such that sparseness is enforced by inhibition alone (not thresholds), then variable inhibition can equalize KC activity without becoming negative (Fig. 5G). However, in this case, the coding level without inhibition was 99.7% (Fig.5G), which is not observed experimentally [25]. Even allowing a coding level without inhibition of 50%, equalizing KC activity still requires some APL-KC inputs to be negative (Fig. 5G). Overall, these results suggest that tuning inhibitory weights cannot compensate on its own for variability in other KC parameters. More likely, the system optimizes multiple parameters at once (see Fig. 7 and Discussion).

To better understand why equalizing average activity improves performance, we asked whether memory performance can also be rescued by equalizing not KC average activity, but rather KC response probability (equivalent to average activity if KC activity is binarized, i.e., 0 or 1). Equalizing response probability (as opposed to average activity) by tuning KC spiking thresholds has been shown to improve separation of KC odor representations in a different computational model [34]. However, in our model, this technique (tuning thresholds to equalize KC response probability) produced worse classification performance and lower dimensionality compared to tuning thresholds to equalize KC average activity (Fig. S4A,B), though still better than the random model (compare Fig. S4 to Fig. 5). This result can be understood by considering that dimensionality of neuronal activity is maximized when variance along all dimensions is equal (Fig. 3) [40], but equalizing KC response probability still allows KCs to have unequal average activity (one KC’s supra-threshold activity might be higher than another’s), which would cause KCs to differ between each other in their variances in activity across odors (a KC’s variance in activity depends on its average activity because its response to most odors is zero).

### Robustness of pre-tuned compensations in new environments with novel odors

Any activity-dependent tuning depends on the model’s context. If a fly tunes its network parameters based on experience in one odor context (e.g., smelling only odors of one chemical family), will it still perform well at classifying odors in a novel environment with different odors (e.g., odors of a different chemical family)? We hypothesized that performance would depend more on tuning context with the activity-dependent compensation mechanisms than the activity-independent mechanism where input weights were picked depending on *N* and *θ* rather than activity.

To test this, we tuned the parameters in our models using only a subset of odors from [30], grouped by chemical class, and then trained and tested the models on odor-reward/punishment associations using the other odors. We took the four chemical classes that had the most odors in the dataset: acids, terpenes, alcohols and esters. For each class, we tuned the model’s parameters on that class and then trained the model to classify odors in the other 3 classes (‘novel’ environment). For matched controls, we trained models that had been tuned on the same 3 classes used for training/testing (‘familiar’ environment). As expected, the three activity-dependent models performed worse in novel environments than familiar environments, while the activity-independent model performed consistently regardless of tuning environment (blue, green and magenta vs. cyan in Fig. 6C). However, in general, tuning odors on one class but training/testing on different classes does not fatally damage the activity-dependent compensation strategies: although performance is worse in novel environments, it remains better than the random model. Thus, activity-dependent compensation is still a good strategy to overcome the pernicious effects of inter-KC variation, even if the compensation environment differs from the classification environment (at least within the range of the odors in [30]).

**Figure 6:**
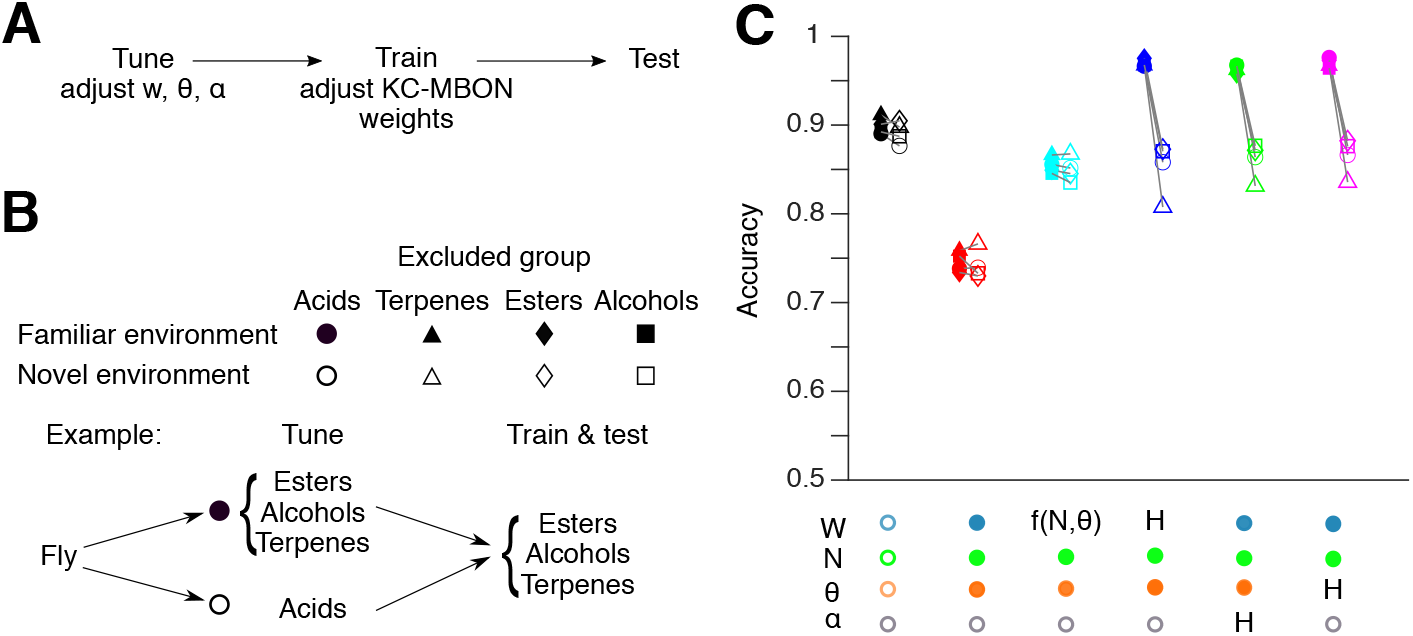
Robustness of pre-tuned compensations with novel odors. (**A**) For each model fly, network parameters are tuned as in Fig. 5, on a subset of odors. At this stage, no rewards or punishments are given, and KC output weights are not modified. Then, the model is trained to classify rewarded and punished odors that are the same as or different from the odors used for tuning. Finally, the model is tested on new noisy variants of the odors used for training. (**B**) Empty symbols (‘novel’ environment): models were tuned on odors from one chemical group (*Gi*: acids - circles, terpenes - triangles, esters - diamonds, or alcohols - squares), then trained and tested on odors from the other three groups (*G*_*i*≠*j*_). Each empty symbol is paired with a matched control (filled symbols) showing how that model would have fared in a ‘familiar’ environment: a model tuned, trained, and tested all on the same three groups of odors as the matched ‘novel’ model was trained and tested on (*G*_*i*≠*j*_). (**C**) Models with activity-dependent compensation (blue, magenta, green) performed worse in novel environment than familiar environments (matching indicated by connecting lines). In contrast, models with no compensation (black, red), or compensation based on network parameters alone rather than activity (cyan), performed similarly in novel and familiar environments. Mean of 20 model instantiations, where each instantiation received a different permutation of odors (see Methods). Annotations below graph indicate whether parameters were fixed (empty circle), variable (filled circle), or variable following a compensation rule (‘H’ for homeostatic tuning, *f* (*N, θ*) for parametric tuning). Differences between novel and familiar environments, *p* < 0.05, Wilcoxon signed-rank test, except for: homogeneous model (black), esters; compensation by parametric tuning (cyan), acids, terpenes, esters (full statistics in Table S1).

### Connectome reveals compensatory variation of input strength and numbers

Our proposed compensatory mechanisms predict correlations between the key model parameters. Excitatory weights (*w*) should be inversely correlated to number of PNs per KC (*N*) where *w* is tuned to compensate for variable *N* and *θ* (Fig. 7B) or where *w* is tuned to equalize KC activity (Fig. 7C). Meanwhile, inhibitory weights (*α*) should be positively correlated to the sum of excitatory weights (Σ*w*, or 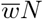, where 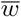 is the mean *w* per KC) where inhibitory weights are tuned to equalize KC activity (Fig. 7D). Such correlations have been observed in larvae [43], but they have not yet been analyzed in the adult mushroom body.

**Figure 7:**
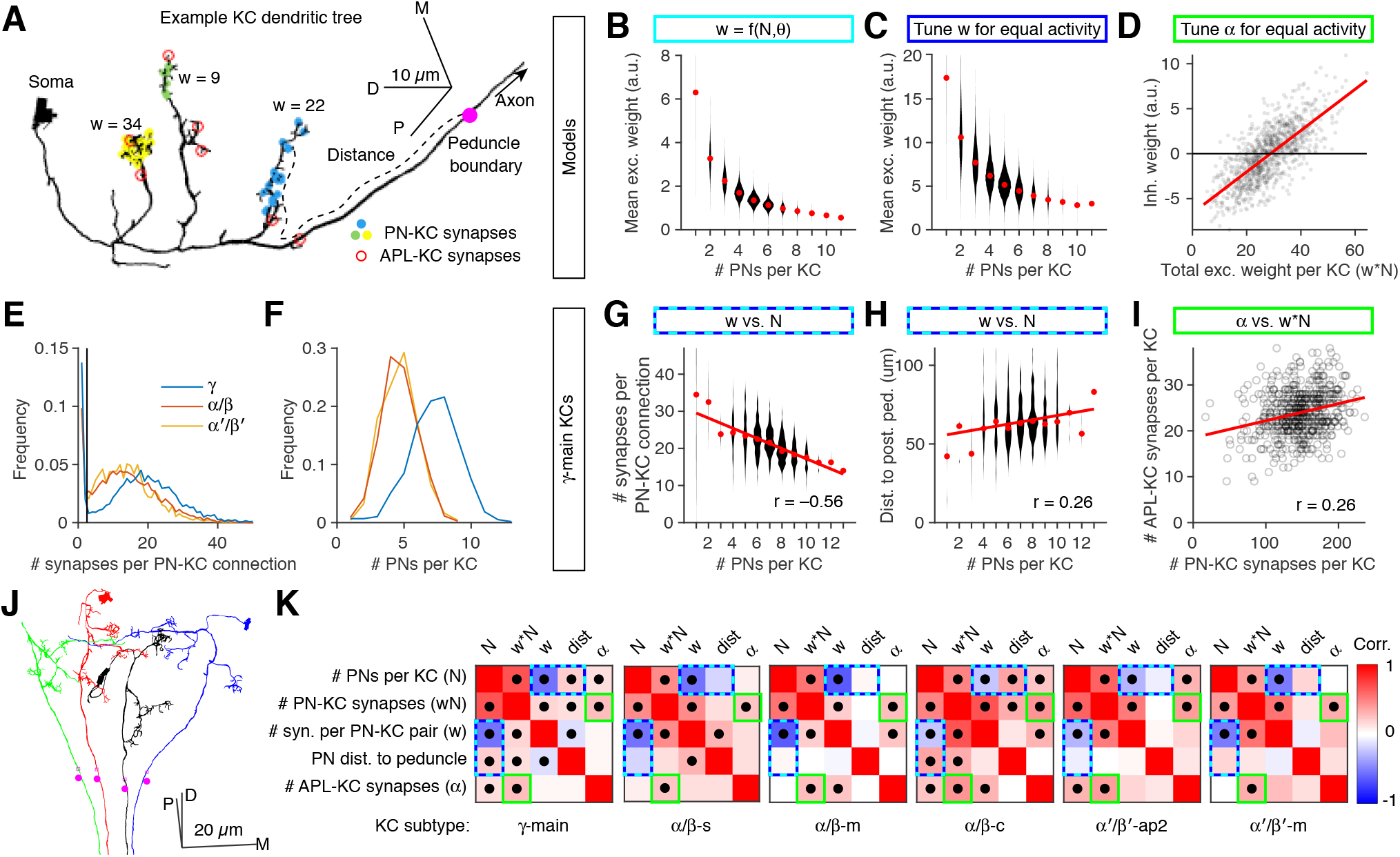
Connectome analysis reveals compensatory variation in excitatory and inhibitory input strengths. **(A)** Example *αβ*-c KC (bodyId 5901207528) with inputs from 3 PNs (yellow/green/blue dots) and 7 dendritic APL-KC synapses (red circles). The magenta circle shows the posterior boundary of the peduncle. Line widths not to scale. **(B,C)** Mean synaptic weight (*w*) per PN-KC connection is inversely related to the number of input PNs in models that tune input weights given *N* and *θ* **(B)**, or that tune input weights to equalize average activity levels across KCs **(C)**. **(D)** In the model that tunes input inhibitory synaptic weights (*α*) to equalize average activity levels across KCs, inhibitory weights are directly related to the sum of excitatory weights per KC (i.e., *wN*). Note the negative values of *α* (discussed in text). **(E,F)** Probability distributions of the number of synapses per PN-KC connection **(E)** and the number of input PNs per KC (**F**) in the different KCs subtypes (*αβ, γ, α′β′*). (**G**) Mean number of input synapses per PN-KC connection is inversely related to the number of input PNs per KC, in *γ*-main KCs (see Fig. S5 for other KC types). (**H**) Mean distance of PN-KC synapses to the posterior boundary of the peduncle (presumed spike initiation zone) is directly related to the number of input PNs per KC. (**I**) The number of APL-KC synapses per KC is directly related to the total number of PN-KC synapses per KC. (**J**) Four *αβ*-c KCs, one from each neuroblast clone. The posterior boundary of the peduncle (magenta circles) lies where the KC axons begin to converge. (**K**) Grids show Pearson correlation coefficients (*r*) between various KC parameters for all KC subtypes tested (red: positive; blue: negative). Dots indicate *p* < 0.05 (Holm-Bonferroni corrected) (full statistics in Table S1). Colored outlines indicate predictions of models (cyan/blue: models tuning *w* **(G,H)**; green: model tuning *α* **(I)**). Number of KCs for each subtype, left to right: 588, 222, 350, 220, 127, 119. In (B,C,G,H), red dots are medians and the widths of the violin plots represent the number of KCs in each bin. Trend lines in (D,G,H,I) show linear fits to the data. Scale bars in **(A,J)**: D, dorsal, P, posterior, M, medial.

To test these predictions, we analyzed the recently published hemibrain connectome [44, 45], which annotates all synapses between PNs and KCs in the right mushroom body of one fly. The connectome reveals three of our parameters: the number of PN inputs per KC (*N*), the strength of each PN-KC connection (*w*), and the strength of inhibitory inputs (*α*). Although the anatomy does not directly reveal *w* and *α* (which can only be measured electrophysiologically), we used an indirect proxy for synaptic strength: the number of synapses per connection (i.e., number of sites between two neurons where neuron 1 has a T-bar and neuron 2 has a postsynaptic density, counted by machine vision; Fig. 7A). It seems reasonable to presume that, all else being equal, connections with more synapses are stronger. Indeed, in the *Drosophila* antennal lobe, when comparing connections from ORNs to ipsilateral PNs vs. contralateral PNs, ipsilateral connections are both stronger [46] and have more synapses per connection [47]. Moreover, synaptic counts approximate synaptic contact area throughout the larval *Drosophila* nervous system [48] and synaptic area approximates EPSP amplitude in mammalian cortex [49].

Therefore, to test if mean *w* and *N* are inversely correlated across KCs, we asked if the number of PN inputs per KC was inversely correlated to the number of synapses per PN-KC connection. We ignored PN-KC connections with 2 or fewer synapses, because the number of synapses per PN-KC connection formed a bimodal distribution with a trough around 3-4 (Fig. 7E); we presumed that connections with only 1-2 synapses represent annotation errors. We divided KCs into their different subtypes as annotated in the hemibrain [45], because different subtypes have different numbers of PN inputs per KC and different numbers of synapses per PN-KC connection ([28]; Fig. 7E,F, S5). We excluded KCs that receive significant non-olfactory input (*γ*-d, *γ*-t, *αβ*-p, *α′β′*-ap1). In all analyzed subtypes of KCs (*γ*-main, *αβ*-s, -m and -c; *α′β′*-ap2 and -m), the number of PN inputs per KC (*N*) was inversely correlated to the mean number of synapses per PN-KC connection, averaged across the PN inputs onto a KC (proxy for 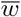) (Fig. 7G,K, S5). Linear regression showed that on average, there were ≈ 6 − 15% fewer input synapses per PN-KC connection 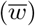, for each additional PN per KC (*N*). This negative correlation meant that the number of total PN-KC synapses per KC increased only sublinearly relative to the number of PN inputs per KC (Fig. S5).

We also tested another anatomical proxy of excitatory synaptic strength. Because KCs sum up synaptic inputs linearly or sublinearly, their dendrites likely lack voltage-gated currents that would amplify inputs, so synaptic input currents likely propagate passively [26]. Therefore, an excitatory input would make a smaller contribution to a KC’s decision to spike the farther away it is from the spike initiation zone [50]. While the spike initiation zone cannot be directly observed in the connectome, the voltage-gated Na^+^ channel *para* and other markers of the axon initial segment (also called the ‘distal axonal segment’) are concentrated at the posterior end of the peduncle, near where axons from KCs derived from the four neuroblast clones converge [51, 52]. This location can be approximated in the connectome as the posterior boundary of the ‘PED(R)’ region of interest (ROI) (magenta dots, Fig. 7A,J). From this point, we measured the distance along each KC’s neurite skeleton (i.e., not the Euclidean distance) to each PN-KC synapse. In the *αβ*-c and *γ*-main KCs (but not other KCs), this distance was positively correlated with the number of PNs per KC (Fig. 7H,K, S5). That is, the more PN inputs a KC has, the farther away the input synapses are from the putative spike initiation zone (and thus the weaker they are likely to be). Intriguingly, of all the KC subtypes, *αβ*-c KCs show the strongest correlation between number of PN inputs and PN-peduncle distance, but the weakest correlation between number of PN inputs and number of synapses per PN-KC connection (Fig. 7K), suggesting that different types of KCs might use different mechanisms to achieve the same compensatory end.

To test if inhibitory and excitatory input are positively correlated across KCs (as predicted in Fig. 7D), we approximated *α* by counting the number of synapses from the APL neuron to every KC in the calyx (the ‘CA(R)’ ROI). In all types of KCs, the more total PN-KC synapses there were per KC, the more calyx APL-KC synapses there were (Fig. 7I,K, S5), indicating that indeed, inhibitory and excitatory synaptic input are correlated.

These results confirm the predictions of our compensatory models. That correlations exist for both excitation and inhibition suggests that the mushroom body tunes more than one parameter simultaneously (thresholds may be tuned as well, but cannot be measured in the connectome). Such multi-parameter optimization likely explains (1) why the correlations in the connectome are not as steep as when only a single parameter is tuned in our models (Fig. 7D-F), and (2) why natural compensatory variation of tuned parameters need not be as wide as the variation of tuned parameters in our models (Fig. 5F).

## Discussion

Here we studied the computational costs and benefits of inter-neuronal variability for associative memory. Using a computational model of the fly mushroom body, we showed that associative memory performance is reduced by experimentally realistic variability among Kenyon cells in parameters that control neuronal excitability (spiking threshold and the number/strength of excitatory inputs). These deficits arise from the reduced separability and dimensionality of odor representations, which arises from unequal activity levels among Kenyon cells. However, memory performance can be rescued by using variability along one parameter to compensate for variability along other parameters, thereby equalizing average activity among KCs. These compensatory models predicted that certain KC features would be correlated with each other, and these predictions were borne out in the hemibrain connectome. In short, we showed (1) the computational benefits of compensatory variation, (2) multiple mechanisms by which such compensation can occur, and (3) anatomical evidence that such compensation does, in fact, occur.

Note that when we say “equalizing KC activity”, we do not mean that all KCs should respond the same to a given odor. Rather, in each responding uniquely to different odors (due to their unique combinations of inputs from different PNs), they should keep their *average* activity levels the same. That is, while KCs’ odor responses should be heterogeneous, their average activity should be homogeneous.

How robust are our connectome analyses? We found correlations between anatomical proxies for the physiological properties predicted to be correlated in our models (i.e., KCs receiving excitation from more PNs should have weaker excitatory inputs, while KCs receiving more overall excitation should also receive more inhibition). In particular, we measured the number of synapses per connection as a proxy for the strength of a connection. As described above, this proxy seems valid based on matching anatomical and electrophysiological data [47–49]. However, other factors affecting synaptic strength (receptor expression, post-translational modification of receptors, pre-synaptic vesicle release, input resistance, etc.) would not be visible in the connectome. Of course, such factors could further enable compensatory variability (see below), so anatomical proxies may actually underestimate the strength of correlations between physiological properties.

We also used the distance between PN-KC synapses and the peduncle as a proxy for the passive decay of synaptic currents as they travel to the spike initiation zone. In the absence of detailed compartmental models of KCs, it is hard to predict exactly how much increased distance would reduce the effective strength of synaptic inputs, but it is plausible to assume that signals decay monotonically with distance. Note that calcium signals are often entirely restricted to one dendritic claw [26, 53]. Another caveat is that the posterior boundary of the peduncle is only an estimate (though a plausible one: [51, 52]) of the location of the spike initiation zone. However, inaccurate locations should only produce fictitious correlations for Fig. 7J and S5H if the error is correlated with the number of PN-KC synapses per KC (and only in *αβ*-c and *γ*-main KCs, not other KCs), which seems unlikely.

Our work is consistent with prior work, both theoretical and experimental, showing that compensatory variability can maintain consistent network behavior [1–11, 54, 55]. However, to our knowledge, we are the first to analyze the computational benefits of equalizing activity levels across neurons in a population (as opposed to across individual animals or over time). A recent pre-print showed that equalizing response probabilities among KCs reduces memory generalization [34], but we showed that equalizing average activity outperforms equalizing response probabilities (Fig. S4), because only the former equalizes variance in activity among KCs to maximize dimensionality. Another model of the mushroom body used compensatory inhibition, in which the strength of inhibition onto each KC was proportional to its average excitation [31], similar to our inhibitory plasticity model (Fig. 5A2). However, the previous work did not analyze the specific benefits from the compensatory variation; it also set the inhibition strong enough that average net excitation was zero, whereas we show that when inhibition is constrained to be only strong enough to reduce KC activity by ≈ half (consistent with experimental data: [25]), inhibition alone cannot realistically equalize KC activity (Fig. 5G). In addition, there is experimental support for our models’ predictions that KCs with more PN inputs would have weaker excitatory inputs: when predicting whether calcium influxes in individual claws would add up to cause a supra-threshold response in the whole KC, the most accurate prediction came from dividing the sum of claw responses by the log of the number of claws [53]. However, the functional benefits of this result only become clear with our computational models. Finally, the larval mushroom body shows a similar relationship between number and strength of PN-KC connections: the more PN inputs a KC has, the fewer synapses per PN-KC connection [43]; however, again, the larval work did not analyze the computational benefits of this correlation.

We modeled two forms of compensation: direct correlations between neuronal parameters (Fig. 5A1) and activity-dependent homeostasis (Fig. 5A2-4). Both forms improve performance and predict observed correlations in the connectome. We cannot directly resolve which mechanism explains the connectome correlations, but can speculate by comparing whether key parameters are correlated with the number of PN inputs (*N*) but not total number of PN-KC synapses 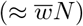, which would suggest a mechanism based on dendritic morphology rather than activity, or vice versa (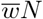 but not *N*), which would suggest the opposite. Where PN-peduncle distance shows significant correlations, it is correlated with both number of PN inputs and total number of PN-KC synapses, suggesting that either mechanism is possible (Fig. 7). Conversely, the number of APL synapses (≈ *α*) is more strongly correlated with the total number of PN-KC synapses than with the number of PN inputs, which is more consistent with activity-dependent tuning than parametric tuning. On the other hand, it may be that *α* is weakly directly tuned to both 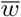 and *N* and thus more strongly tuned to the combination, 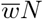.

Certainly, activity-dependent mechanisms are plausible, as KCs regulate their own activity homeostatically in response to perturbations in activity [56]. Indeed, different KC subtypes use different combinations of mechanisms for homeostatic plasticity [56], consistent with the different correlations observed in the connectome for different KC subtypes. Our activity-dependent models lend themselves to straightforward biological interpretations. Excitatory or inhibitory synaptic weights could be tuned by activity-dependent regulation of number of synapses per connection or expression/localization of receptors or other post-synaptic machinery. Spiking thresholds could be tuned by altering voltage-gated ion conductances or moving/resizing the spike initiation zone [52, 57].

On the other hand, KCs are not infinitely flexible in homeostatic regulation; for example, complete blockade of inhibition causes the same increase in KC activity regardless of whether the blockade is acute (16 - 24 h) or constitutive (throughout life) [56]. This apparent lack of activity-dependent down-regulation of excitation suggests that activity-independent mechanisms might contribute to compensatory variation in KCs, as occurs for ion conductances in lobster stomatogastric ganglion neurons [8, 9]. For example, the inverse correlation of *w* and *N* arises from the fact that the number of PN-KC synapses per KC increases only sublinearly with increasing numbers of claws (i.e., PN inputs) (Fig. S5H). Perhaps a metabolic or gene regulatory constraint prevents claws from recruiting postsynaptic machinery in linear proportion to their number. (Interestingly, this suppression is stronger in larvae, where the number of PN-KC synapses per KC is actually constant relative to the number of claws: [43].) Meanwhile, the correlation between number of inhibitory synapses and number of excitatory synapses might be explained if excitatory and inhibitory synapses share bottleneck synaptogenesis regulators on the post-synaptic side. Although activity-dependent compensation produced superior performance in our model compared to activity-independent compensation thanks to its more effective equalization of KC average activity (Fig. 5) (most likely because it takes into account the unequal activity of different PNs), activity-dependent mechanisms suffered when the model network switched to a novel odor environment (Fig. 6). Given that it is desirable for even a newly eclosed fly to learn well, and for flies to learn to discriminate arbitrary novel odors, activity-independent mechanisms for compensatory variation may be more effective in nature.

Compensatory variability to equalize activity across neurons could also occur in other systems. The vertebrate cerebellum has an analogous architecture to the insect mushroom body; cerebellar granule cells are strikingly similar to Kenyon cells in their circuit anatomy, proposed role in ‘expansion recoding’ for improved memory, and even signaling pathways for synaptic plasticity [21, 40, 58–61]. Whereas cortical neurons’ average spontaneous firing rates vary over several orders of magnitude [62], granule cells are, like Kenyon cells, mostly silent at rest, and it is plausible that their average activity levels might be similar (while maintaining distinct responses to different stimuli) [63]. Granule cell input synapses undergo homeostatic plasticity [64], while compartmental models suggest that differences in granule cells’ dendritic morphology would affect their activity levels, an effect attenuated by inhibition [65], raising the possibility that granule cells may also modulate inter-neuronal variability through activity-dependent mechanisms. Future experiments may test whether compensatory variability occurs in, and improves the function of, the cerebellum or other brain circuits. Finally, activity-dependent compensation may provide useful techniques for machine learning. For example, we found that performance of a reservoir computing network could be improved if thresholds of individual neurons are initialized to achieve a particular activity probability given the distribution of input activities [66].

## Methods

### Modelling KC activity

PN activity was simulated using the odor responses of 24 olfactory receptors [30], passed through an equation proposed by [29]. For an ORN and PN innervating the *i*th glomerulus, their responses to the *k*th odor can be described using 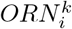 (ORN activity) and 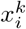 (PN activity):

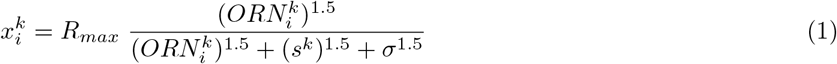

where 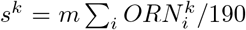, *m* = 10.63, representing the gain of lateral inhibition in the antennal lobe, *R_max_* = 165, representing the maximum PN response, and *σ* = 12, representing the non-linearity of the ORN-PN response function. We added noise to PN activity using:

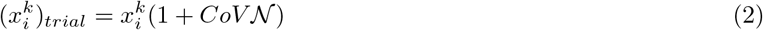

where *CoV* is the coefficient of variation of PN activity across trials taken from Fig. 2E of [35] and 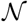 is a random sample drawn from a Gaussian distribution with mean 0 and standard deviation 1. To increase the number of stimuli beyond the 110 recorded odors in [30], we generated odor responses in which the activity of each PN was randomly sampled from that PN’s activity across the 110 odors used in [30], i.e., 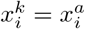 where *k* = 1…*K*, *K* being the number of simulated odors, and *a* is randomly sampled from integers from 1 to 110 for each PN and each odor.

We modeled 2000 KCs. The *j*th KC received input from a randomly selected set of *N_j_* PNs, where *N_j_* was either fixed at 6 or sampled from a Gaussian distribution with mean 6 and standard deviation 1.76 (integer values only), based on experimental measurements from 200 KCs [28]. Although more recent results show that PN-KC connectivity is not entirely random, as KCs that receive inputs from a certain group of food-odor-responsive glomeruli are slightly more likely to receive other inputs from that same group [45, 67], we judged that attempting to model this non-randomness would not add to the realism of our model given that we modeled only 24 (out of ≈50) glomeruli.

The connection from the *i*th PN to the *j*th KC had strength *w_ji_*, which was 0 for non-connected neurons, and for connected neurons was either fixed at 1, sampled from a log-normal distribution (*μ* = −0.0507 and *σ* = 0.3527, based on [27]), or tuned by one of the methods described below. KCs received inhibition from APL (modeled as pseudo-feedforward for simplicity), with a gain that was either constant across all KCs (*α*) or tuned individually as described below (*α_j_*). The KCs’ spiking thresholds *θ_j_* were either constant across all KCs, or sampled randomly from a Gaussian distribution with coefficient of variation 0.26, based on experimental measurements of the difference between spiking threshold and resting potential in 17 KCs [27]. These spiking thresholds were subject to a scaling factor *C_θ_* to achieve the correct average coding level (see below). Thus, the activity of the *j*th KC for the *k*th odor, 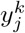, was

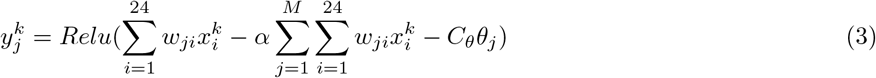

where *M* = 2000 is the number of KCs and *Relu* is a rectified linear unit:

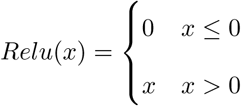

The coding level, or fraction of KCs active for each odor, averaged across odors, was defined as:

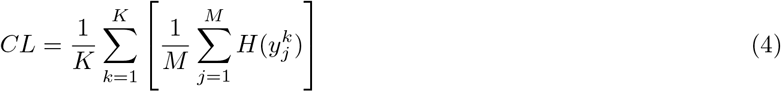

where *K* and *M* are the number of odors and KCs, respectively and *H*(*x*) is the Heaviside function:

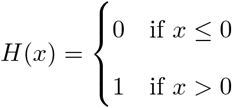

Experimental data suggest that coding level is around 0.1 normally, and approximately double that (0.2) when inhibition is blocked [25]. To match these constraints, we minimized this error function with respect to *C_θ_* (thus preserving the coefficient of variation of thresholds across KCs, i.e., *C_θ_θ_j_*):

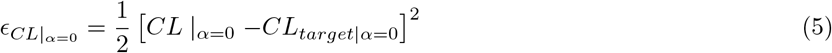

where *CL*_*target*|*α*=0_ = 0.2 and we minimized this error function with respect to *α*:

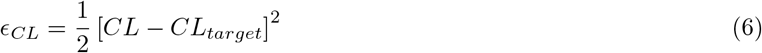

where *CL_target_* = 0.1.

We tuned *C_θ_* and *α* using gradient optimization, using the update equations:

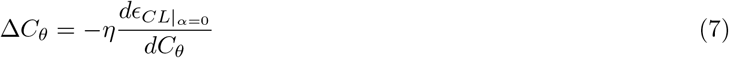

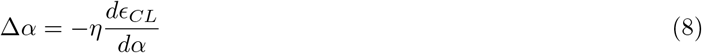

To derive the update rule for Δ*C_θ_*, we differentiate (5) with respect to *C_θ_*:

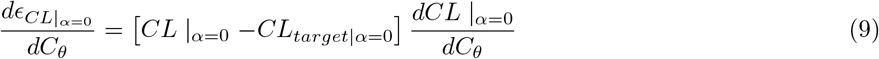

To differentiate *CL* with respect to *C_θ_*, we need to replace the discontinuous Heaviside function with a continuous approximation. Similar to [68] a sigmoid function approximates a Heaviside at the limit *σ* → 0,

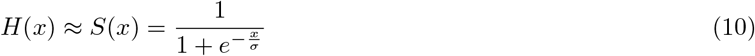

Hence, assuming *σ* = 1, we can define the coding level as:

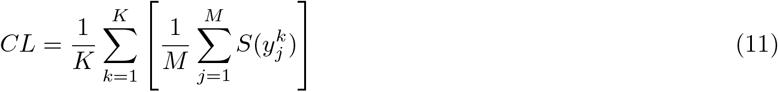

Given the derivative of a sigmoid is:

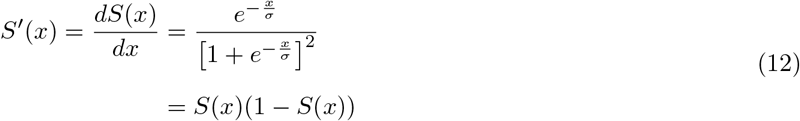

Thus,

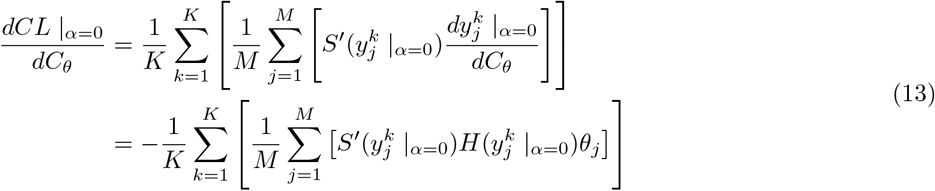

combining (9) and (13), and plugging in (7) we can get the update equation for *C_θ_* as

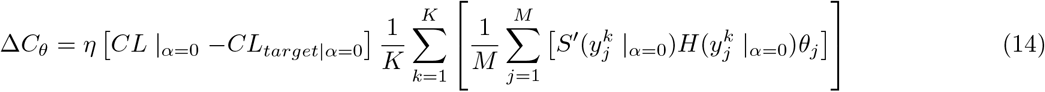

For simplicity, this can be re-written using the average operator notation 〈〉 across odors (indexed by *k*) and KCs (indexed by *j*),

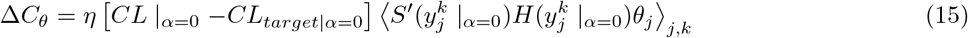

Similarly, for Δ*α* we differentiate (6) with respect to *α*,

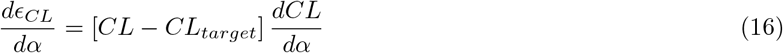

Similarly,

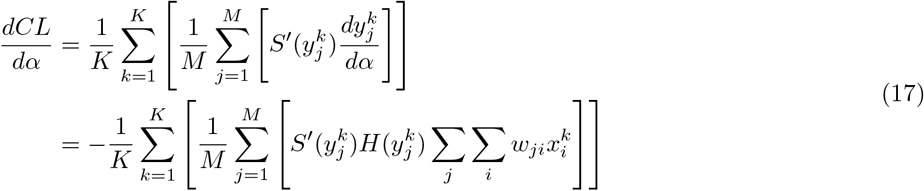

combining (16) with (17) then putting in (8),

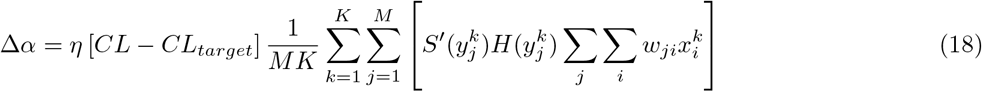

and using the 〈〉 notation:

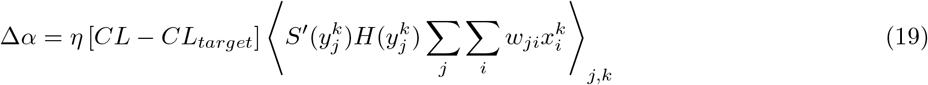

These update equations were used to adjust values of *θ* and *α* in any random instantiation of the fly’s network to match the experimentally observed coding levels. Note that because the update equation for *α* is the same for all *j*, the same equation applies when *α_j_* is tuned for each KC (see below).

### Modelling olfactory associative learning

Learning occurred through synaptic depression at the output synapse from KCs onto MBONs according to this exponential decay rule:

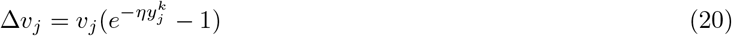

where *v_j_* is the synaptic weight between the *j*th KC and the MBON of the ‘wrong’ valence and *η* is the learning rate. Thus, KCs active for a punished odor weaken their synapses to the approach MBON while KCs active for the rewarded odor weaken their synapses to the avoid MBON. This can be seen as the model fly learning from ‘mistakes’ during its training phase [69, 70].

The behavior of the fly was determined by a softmax equation:

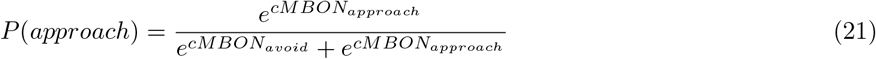

where the constant *c* governs how probabilistic or deterministic the decision-making is. At high *c*, the model approaches a completely deterministic model where the fly will approach the odor 100% of the time whenever the approach MBON’s activity is higher than the avoid MBON’s activity; at very low *c*, the model approaches random chance; in between, the fly’s behavior is probabilistic but biased by the imbalance between the activity of the two MBONs.

We trained the model on 15 noisy trials of the odors (no repetitions) and tested it on 15 unseen noisy trials of the same odors, and calculated the accuracy as the fraction of trials in which the model behaved correctly (i.e., avoided punished odors and approached rewarded odors).

### Metrics for evaluating Kenyon cell odor representations

Angular distance between two vectors *A* and *B* was calculated using:

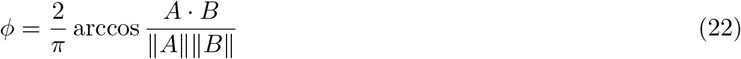

Dimensionality was calculated according to the equation in [40]:

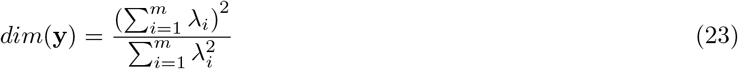

where *λ_i_* are the eigenvalues of the covariance matrix of **y**. Whereas Litwin-Kumar et al. calculated dimensionality analytically given inputs with defined distributions, we calculated it numerically given simulated PN inputs. Because dimensionality cannot be accurately calculated with a small number of inputs, we simulated KC activity for 1000 input odors for dimensionality calculations.

Sparseness was calculated according to [25, 41]. Using the notation of this paper, the lifetime sparseness of the *j*th KC for a set of *K* odors is:

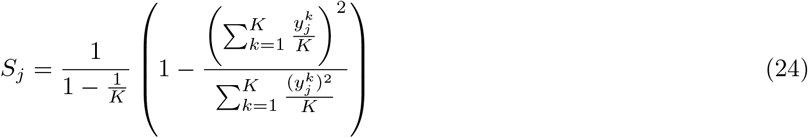

If a cell is completely silent, firing to no stimuli, 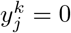 for all *k* and sparseness is undefined due to division by zero.

We used the Davies-Bouldin Index (DBI; [39]) to measure the degree of separation between clusters of the KCs responses for two odors, or between the clusters of the rewarded odors responses versus the punished odors responses. The DBI measures the ratio between the within-cluster variance and the inter-cluster distance. Let clusters *C*_1_ and *C*_2_ consist of sets of *A* and *B N* -dimensional data points, *X* = {*x*_1_*, x*_2_, …*x_A_*} and *Y* = {*y*_1_*, y*_2_, …*y_B_*}, respectively. The DBI is defined as:

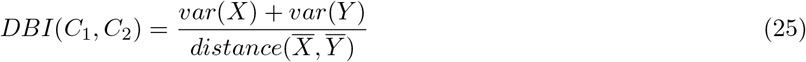

where 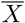 and 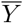 are the centroids of clusters *C*_1_ and *C*_2_, and *distance*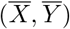 is the Euclidean distance between the two, while *var*(*X*) and *var*(*Y*) are the within-class variances, such that,

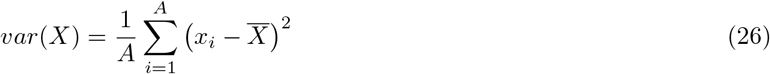

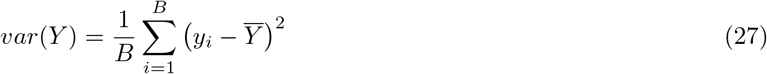

High DBI indicates poor separation (more overlap) between clusters *C*_1_ and *C*_2_, due to either high within-cluster variance or low inter-cluster distance.

### Models for compensatory variability

#### Parametric tuning of excitatory input weights

We approximated the probability distribution of PN-KC synaptic weights (*w*) using the distribution of amplitudes of spontaneous excitatory post-synaptic potentials (mini-EPSPs) in KCs, measured by [27]. This experimental distribution was approximately log-normal, as has been described for cortical synapses [62, 71], so we modeled *w* as following a log-normal distribution. We simulated values of *w* such that the overall distribution of *w* would follow this log-normal distribution, yet individual KCs would sample *w* from different log-normal distributions depending on *N* and *θ*, such that KCs with lower *N* or higher *θ* would have higher *w*, i.e., sampling from a log-normal distribution shifted to the right (Fig. 5A1).

The probability of PN-to-KC synaptic weights could be estimated from the probability summation rule,

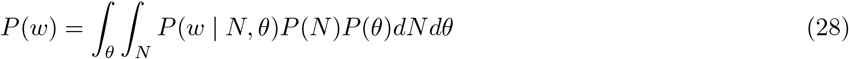

where *P* (*w* | *N, θ*) is the conditional probability distribution of the input synaptic weights for a KC that has *N* claws and spiking threshold *θ*, sampled from probability distributions *P* (*N*) and *P* (*θ*), respectively. We approximated *P* (*N*) and *P* (*θ*) as the Gaussian distributions described above (see Fig. 2), and we approximated integration over *θ* as summation at small intervals (Δ*θ* = 2.5).

We modeled the constituent conditional probability distributions *P* (*w* | *N, θ*) as also being log-normal, based on previous studies which approximate the sum of log-normal distributions as another log-normal variable by matching the first two moments of the power sum and its individual log-normal contributors [72–74]. This approximation holds in our case (the Kullback-Leibler Divergence metric (KLD) converged to less than 0.001).

To get the posterior lognormal distributions *P* (*w* | *N, θ*), we minimized the distance metric Kullback-Leibler Divergence (KLD) between *P* (*w*) and ∫_*θ*_ ∫_*N*_ *P* (*w* | *N, θ*)*P* (*N*)*P*(*θ*)*dNdθ*. To implement compensatory tuning in these conditional probabilities, such that a KC with fewer inputs (lower *N*) or higher spiking threshold (higher *θ*) would have stronger inputs (higher median *w*), we parameterized the medians 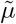 of each conditional distribution in *N* and *θ* as:

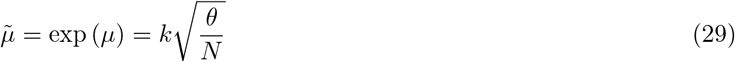

Thus,

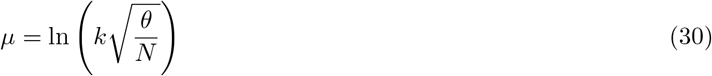

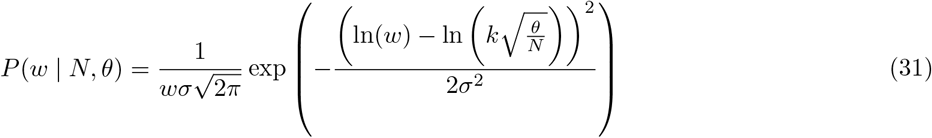

We used gradient descent optimization to find the values of *σ* and *k* in Eq. 31 that would minimize the fitting error:

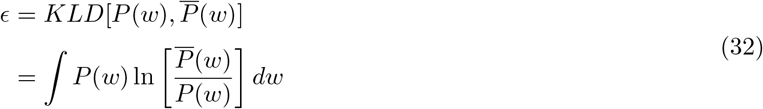

where

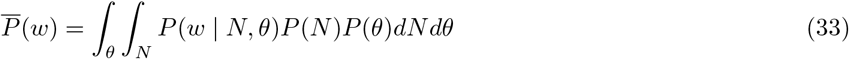

First, we found the optimal *σ* by gradient optimisation:

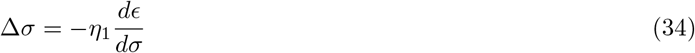

The derivative of the fitting error with respect to *σ* is:

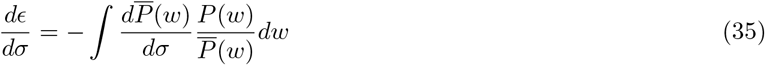

with,

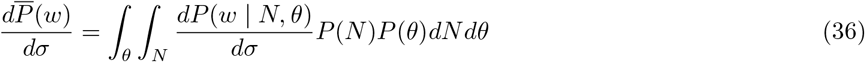

where 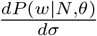 is:

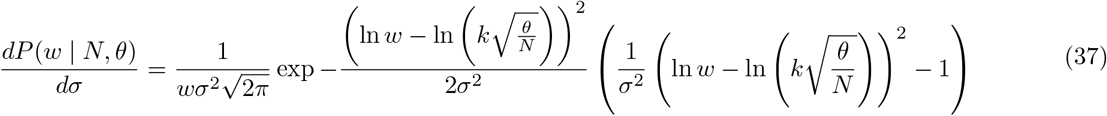

Similarly for *k*,

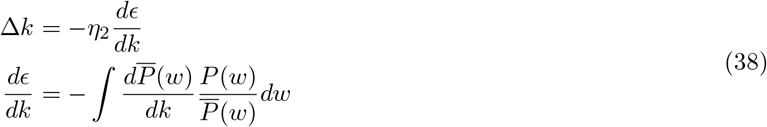

such that,

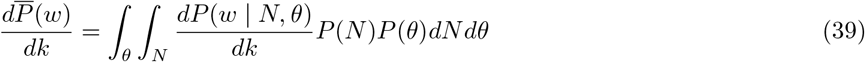

with 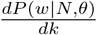 given by:

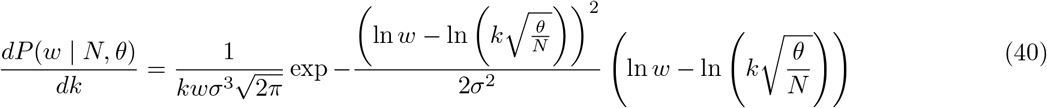

Starting from arbitrary values for *k* and *σ* and using small learning rates *η*_1_ and *η*_2_, at each iteration, the gradient descent algorithm alternated between using *σ* to update *k* and using *k* to update *σ*. We stopped the gradient descent (i.e., the algorithm converged) at *ϵ* < 0.001.

### Tuning KC input excitatory weights to equalize KC activity

In this model, we reduce the high variance in KCs’ average activity levels by tuning their input synaptic weights, such that each *j*th KC adjusts its input synaptic weights (*w_ji_*) to make its average activity level 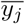 reach a certain desired level *A*_0_. We initially analyzed this problem using an error function:

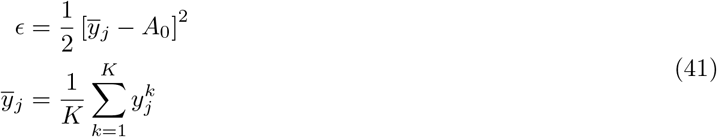

where 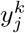 is the *j*th KC’s response to the *k*th odor calculated as in equation (3) and *K* is the number of odors. Finding the weights to minimize the error in (41) can be found by gradient optimisation,

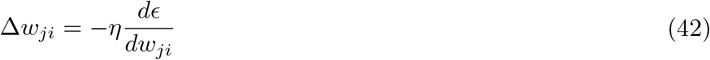

with,

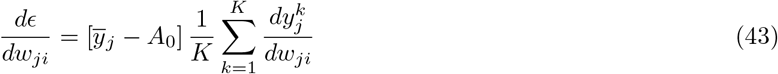

Taking the derivative of 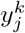 w.r.t. *w_ji_* yields:

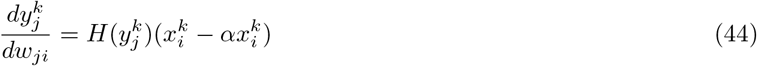

Plugging (44) in (43) gives:

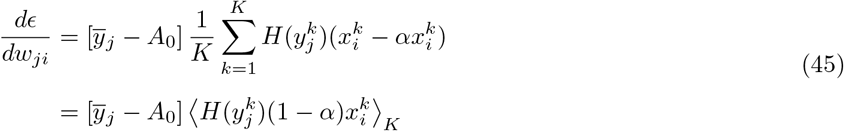

Hence, *w_ji_* will be updated as follows:

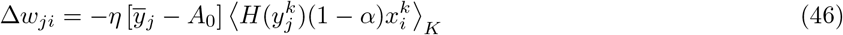

The equation above means that a KC with an average activity 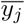 higher (lower) than *A*_0_ will scale down (up) its input synaptic weights, *w_ji_*, proportional to both the difference 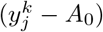 and the average input activity from the *i*th PN. Note that in this derivation a KC must have non-zero average activity, i.e., 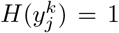 for at least one odor, for its weights to be updated. We believe such a rule would be biologically implausible, as there should not be a discontinuity between a silent KC and a nearly silent KC. To allow totally silent KCs (which have only subthreshold activity) to update their weights in the same way as active KCs, we heuristically apply the following rule:

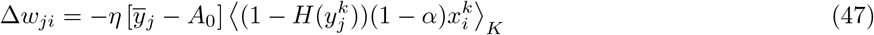

Adding (46) and (47) we obtain:

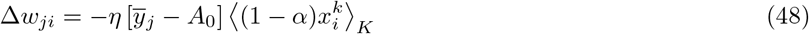

The rule has a fixed point 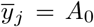 since 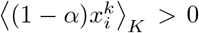. Note that we apply the constraint *w_ji_* ≥ 0. How updates for *w_ji_* = 0 are treated depends on the reason why *w_ji_* = 0: if the *i*th PN and *j*th KC are not connected, then the update is not applied. But if they were originally connected and the update rule pushed *w_ji_* to zero, the update rule will continue to be applied.

To test whether performance is affected by adding the heuristic term to allow silent KCs to update their weights, we compared the performance using update rule Eq. (46) vs. (48). The rule without the heuristic performed significantly worse and had lower dimensionality than the rule with the added heuristic for activating silent KCs (Fig. S3A,B). This means that a formally derived update rule for *w* was not enough, since it would not equalize activity for all KCs (silent KCs will remain silent) and would not enhance the population coding as in the heuristic rule.

We further noted that Eq. (48) contains a factor 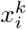 meaning that the update to *w_ji_* depends on the average input activity from the *i*th PN. As this rule makes the biological interpretation more complex (the synaptic update depends on both pre- and post-synaptic activity), we also tested a simplified rule where synaptic changes depend only on the average KC activity:

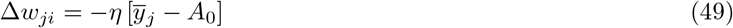

This simplification did not affect memory performance or the tuned distribution of weights (Fig. S3A1,A2,D), but it improved the KCs’ dimensionality (Fig. S3B) and the robustness of the model to novel odor environments (Fig. S3E). This improvement in the model robustness might be because including the extra factor 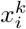 in the learning rule caused the model to be overfitted to the tuning environment. Therefore, we used Eq. (49) for the results presented in the main figures, as it is simpler and produces better performance, despite not being formally derived from an error function. As with Eq. (48), this update rule has a fixed point 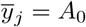.

### Tuning KC input inhibitory weights to equalize average KC activity

In this model, we model each KC as adjusting its individual input inhibitory synaptic weights from APL, to match its average activity level 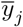 to a certain desired level *A*_0_. We minimize the error function in Eq. (41) by adjusting *α_j_* instead of *w_ji_*:

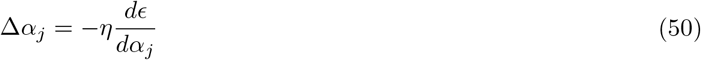

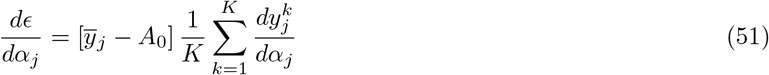

Differentiating 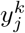 with respect to *α_j_* yields

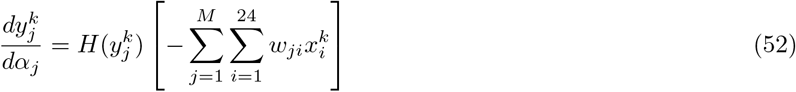

Plugging (63) in (51) gives,

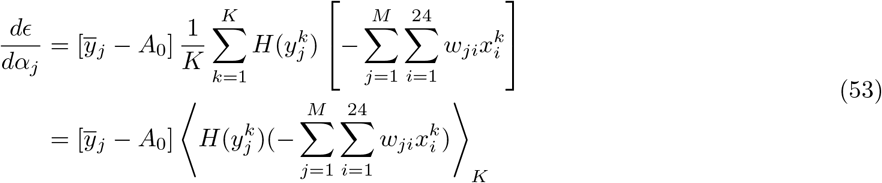

Therefore,

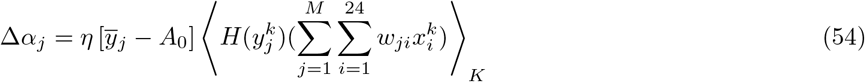

Similar to the previous section, we assume that weight changes for silent neurons happen in the same way as for active neurons:

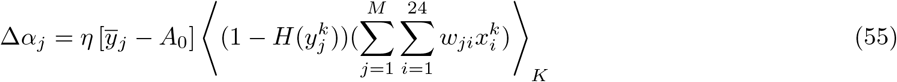

Adding (54) and (55) we obtain the inhibitory plasticity rule allowing KCs to achieve equal average activity:

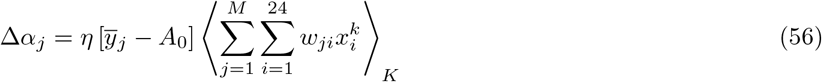

Given that 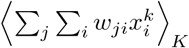 is a constant as *w_ji_* is not updated in this model, this term can be subsumed into the learning rate, so this equation reduces to:

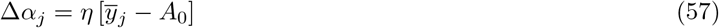

Besides the homeostatic tuning of the APL inhibitory feedback values, these individual values of *α_j_* also have to satisfy the sparsity constraint in Eq. (5). Therefore, the learning rule for these inhibitory weights requires simultaneously optimizing both error functions, Eq. (5) and (41). Thus combining Eq. (56) and the derivative of the sparsity constraint (CL=10%) with respect to each value of *α_j_*,

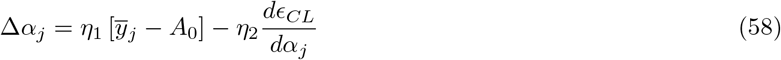

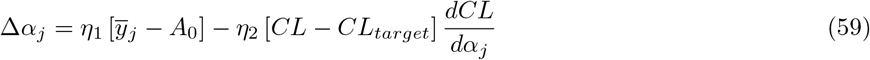

where

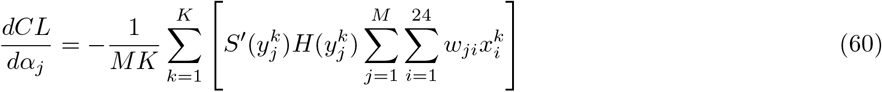

Combining (59) with (60),

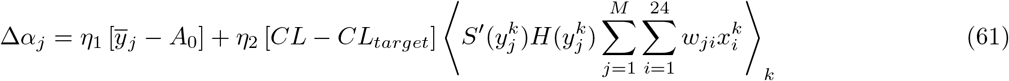

We tested re-parameterizing *α_j_* into *C_α_α_j_* where *C_α_* is tuned across all KCs to adjust coding level while *α_j_* is tuned individually to equalize KC activity levels, but this had no effect on memory performance, so we kept the simpler model formulation.

### Tuning KC spiking thresholds to equalize average KC activity

In this compensatory technique, we tune individual KCs’ spiking thresholds *θ_j_* to achieve equal average activity across the KC population. Starting with arbitrary initial values, each KC adjusts its spiking threshold so its average activity across *K* odors reaches a target level, *A*_0_, by minimizing the error in average activity as in Eq. (41) by gradient optimization:

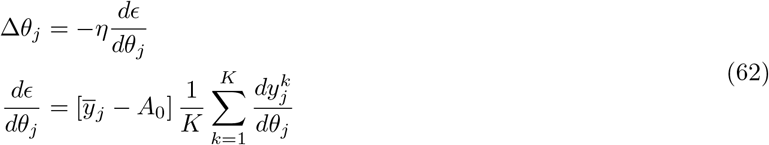

Differentiating 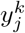, the expression in Eq. (3), with respect to *θ_j_* yields

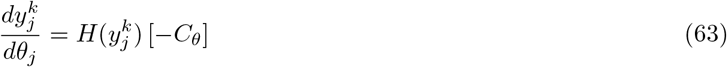

Plugging (63) in (62) gives,

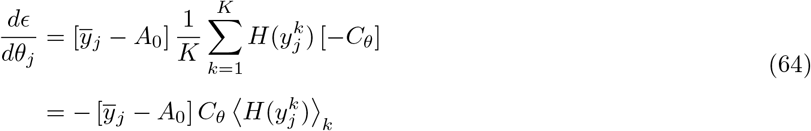

Therefore,

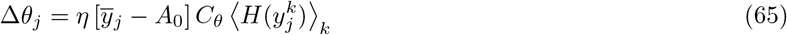

Similar to Eq. (47), we assume that spiking thresholds are updated for silent KCs as well:

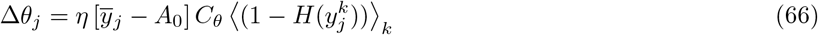

Adding (65) and (66) we obtain the spiking thresholds plasticity rule allowing KCs to achieve equal average activity:

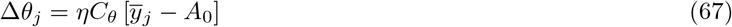

### Tuning spiking thresholds to equalize KCs response probabilities

We tested an alternative strategy to tune *θ* suggested in [34]: to equalize not 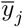 but rather the average response probability of each KC across *K* odors without inhibition, *P_j_*, i.e.:

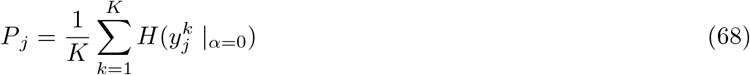

As in Eq. (5), we set this target response probability, 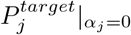, to 0.2 to match experimental findings that blocking inhibition approximately doubles response probability [25]. We minimized the error function:

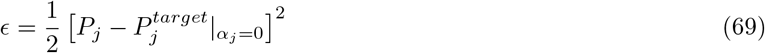

by adjusting *θ_j_* by gradient optimization:

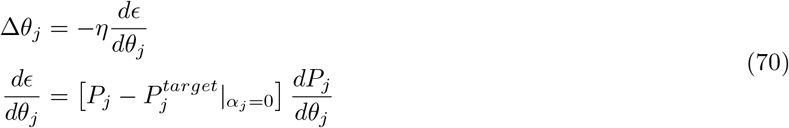

To differentiate *P_j_*, as in Eq. (13), we approximated the discontinuous Heaviside function with a sigmoid:

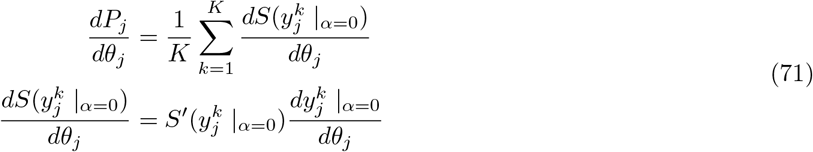

Recalling the formula of 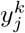 in (3), it follows

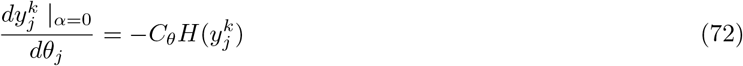

Combining (72) with (71), and plugging in (70),

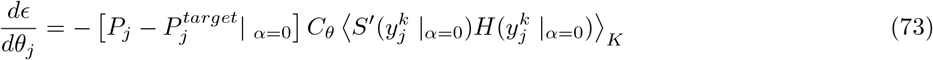

Thus, *θ_j_* values are updated by,

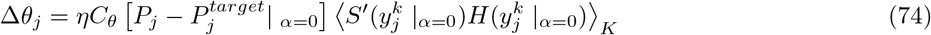

As in Eq. (47), (66) and (55), we can write a symmetric rule for silent KCs:

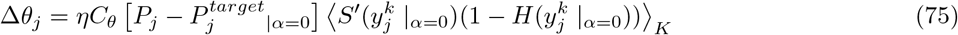

Adding (75) and (74) leads to an activity-dependent update rule for *θ_j_*, given all the incoming input odors:

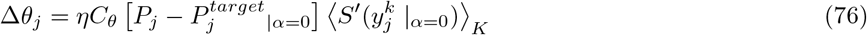

In this model, the sparsity constraint *CL*_*target*|*α*=0_ = 0.2 is satisfied by 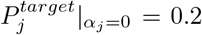, because coding level equals the average of response probabilities across KCs:

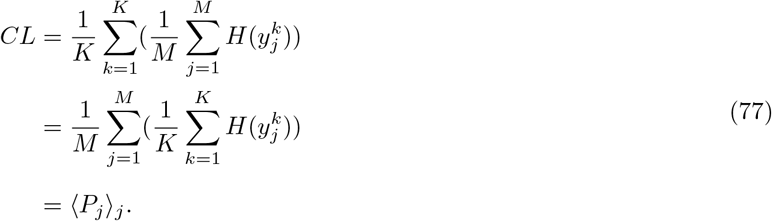

### Optimization of the multiple objective functions

As noted above, homeostatic tuning of *w_ji_*, *θ_j_*, or *α_j_* needs to happen while maintaining the sparsity constraints, Eq. (5) and (6). (It is important to note that the homeostatic update rules are meant to represent a biological process while the sparsity constraints merely fit our model to experimental data and stand in for unknown processes that lead to a coding level of 0.1.) Since these activity-equalizing tunings both depend on and change the network’s sparsity level, we used a sequential optimization approach to optimize each objective function, *O_i_*, at a time. For each *i*, we find the optimal parameters {*P_i_*} minimizing an objective *O_i_*, using the current estimates of the other parameters {*P_j_*} from all the other objectives, {*O_j_*} where *j* ≠ *i*. The algorithm iterates for all *i* to minimise each of the objective functions, until it reaches a global minimum where the errors from all of the objective functions fall below a certain tolerance, *τ_O_*.

Given an initial estimate for *C_θ_*, *α*, *θ_j_* and *w_ji_*, the algorithm goes as follows:

**Algorithm 1:**
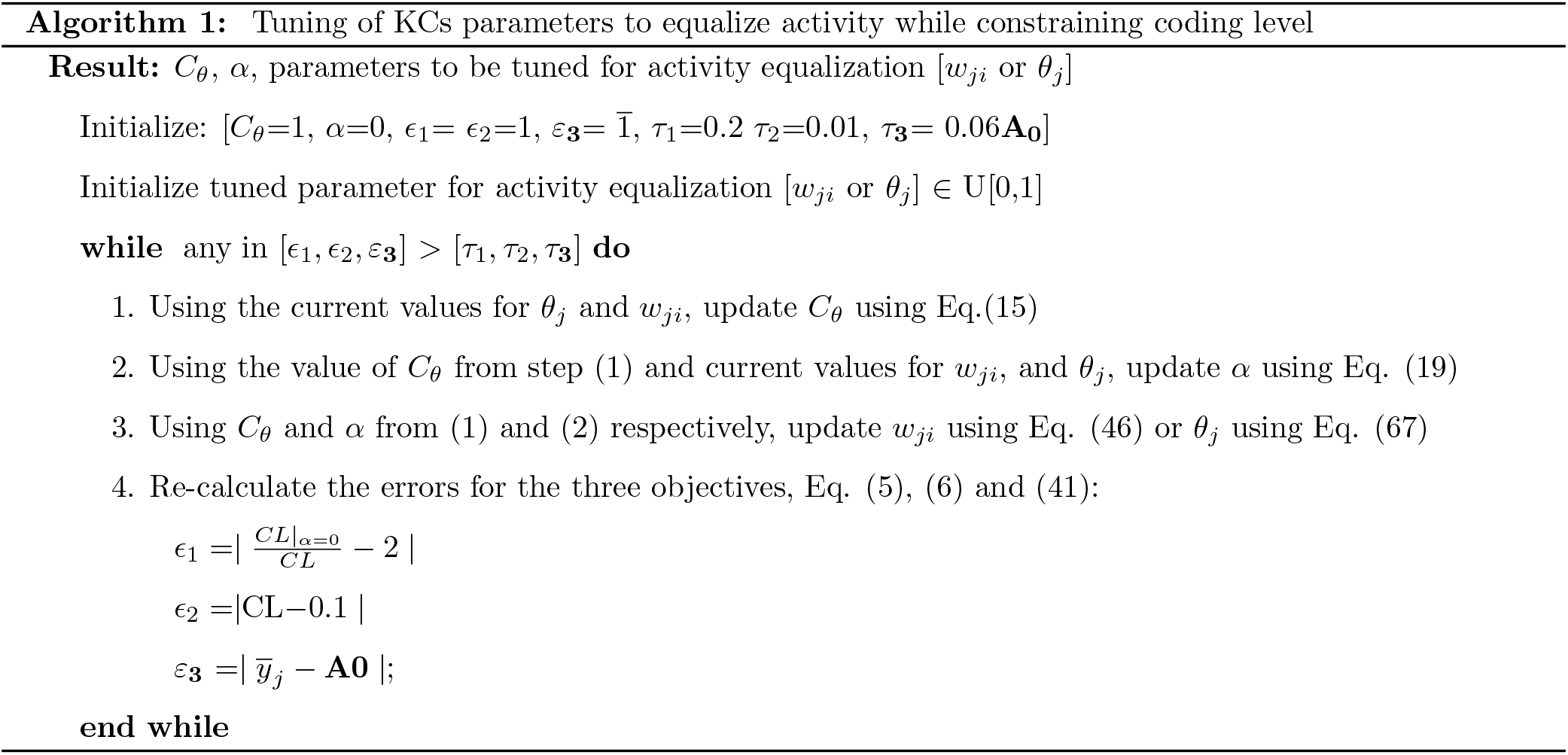
Tuning of KCs parameters to equalize activity while constraining coding level

In our implementation we initialize the parameters to be tuned for activity equalization (*w_ji_*, *θ_j_* or *α_j_*) from a uniform random distribution *U* = [0, 1] (the non-tuned parameters follow the distributions in Fig. 2). In addition, we set the error for the first and second sparsity constraint, Eq. (5) and (6), to be 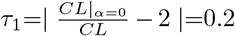, while *τ*_2_= | *CL* − 0.1 |=0.01 respectively. This means allowing the coding level without and with the APL feedback to fall within [1.8*CL* ≤ *CL* |_*α*=0_≤ 2.2*CL*], and [0.09 ≤ *CL* ≤ 0.11] respectively. For the activity equalization objective, the error *ε*_**3**_ is a column vector of size *M*, of the differences between the target average activity value *A*_0_, and the current average activity for each KC, 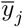. This objective function is satisfied when all the values in the vector *ε*_**3**_ are less than 6% of the target activity.

Note that in the inhibition-tuning model, we tune the same parameter, *α_j_* (a vector of *M* values instead of a constant), to jointly satisfy both the sparsity and the activity-equalization objectives. In this case, step (3) above is removed and step (2) updates *α_j_* using Eq. (61).

In the model where we tune *θ_j_* to equalize response probability rather than average activity (Fig. S4), equalizing response probability without inhibition to 0.2 also solves the coding level constraint (Eq. (77)). Thus, in this case, the algorithm iterates between 2 steps: (1) update *θ_j_* according to Eq. (76), (2) use these values to update *α* according to Eq. (19), as follows,

**Algorithm 2:**
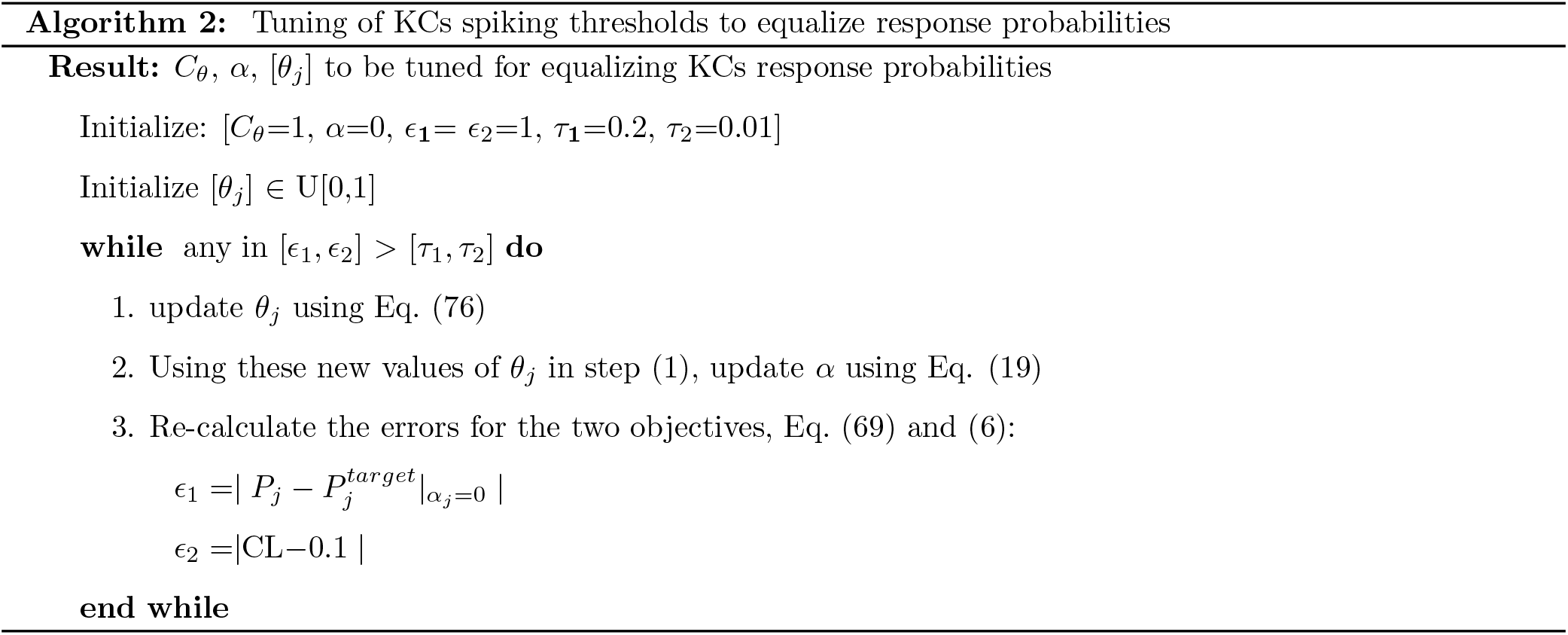
Tuning of KCs spiking thresholds to equalize response probabilities

In our optimization pipeline, there is a potential problem in the models where KC activity is equalized by tuning *α_j_* or *θ_j_*. In these models *w_ji_* is not tuned, so for values of *A*_0_ that are too high relative to values of *w_ji_*, excitation will be too low to reach the high targets given the constraints *C_θ_θ_j_* > 0, *CL* = 0.1 and *CL* |_*α*=0_= 0.2, meaning the algorithm does not converge. (This is not a problem when tuning *w_ji_* because *w_ji_* can go arbitrarily high, whereas thresholds cannot go below zero.) Therefore, *w_ji_* values must be chosen in a sensible range relative to *A*_0_ (keeping in mind that the value of *A*_0_ is arbitrary: see below). Rather than further complicating the objective cost functions by introducing a tunable scaling factor for *w_ji_*, we found that in practice the algorithm converged if *w_ji_* values (starting from a log-normal distribution with *μ* = −0.0507, *σ* = 0.3527) were multiplied by 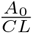 (where *CL* = 0.1). The target activity *A*_0_ is arbitrary because if parameters can be found to satisfy our model constraints (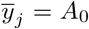, *CL* = 0.1 and *CL* |_*α*=0_= 0.2) for a particular *A*_0_ > 0, then a solution also exists for 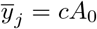 for any *c* > 0, because:

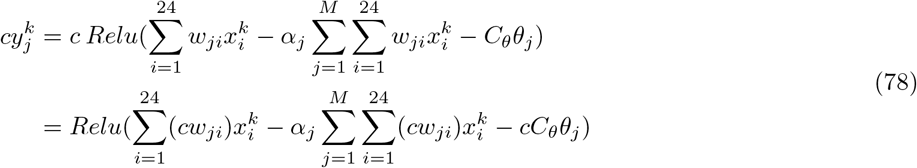

That is, to scale 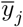 by a factor *c*, one need only scale the parameters *w_ji_* and *C_θ_* by *c*. In other words, only the relative magnitudes of *A*_0_, *w_ji_* and *C_θ_*, not the absolute magnitudes, are meaningful.

### Robustness analysis

Of the 110 odors tested in [30], we took the four chemical classes with the most odors (acids, terpenes, alcohols and esters), so that tuning parameters on a single class would provide a reasonable number of odors (at least 15). Because each class had different numbers of odors, and the memory task is more difficult when more odors need to be classified, we equalized the number of odors in each task by randomly sampling 15 odors from those classes that had more than 15 members (terpenes, 16; alcohols, 18; esters, 24), with a different random sampling for each model instantiation. Because of the small number of odors used for tuning, it was not always possible to equalize the activity of every single KC; in particular, in the threshold-tuning models in a novel environment, we allowed a maximum of 5 KCs to fall outside the ±6% bound on average activity.

### Connectome analysis

KC neurite skeletons and connectivity were downloaded from the hemibrain connectome v. 1.1 [44]. KCs (excluding those that receive significant non-olfactory input) were selected as neurons whose ‘type’ field was `KCg-m, KCab-c, KCab-m, KCab-s, KCa’b’-ap2` or `KCa’b’-m`. PN inputs for a KC were identified as neurons whose ‘type’ field included `adPN, lPN` or `vPN` (NB: some of these, e.g., vPNs, do not project to the mushroom body and so were never counted) and that formed more than 2 synapses with the KC (see Fig. 7B). KCs with truncated skeletons lacking the dendritic tree were excluded. The posterior boundary of the peduncle was the most posterior node in a skeleton annotated as being in the ‘PED(R)’ region of interest (annotations at https://storage.cloud.google.com/hemibrain/v1.1/hemibrain-v1.1-primary-roi-segmentation.tar.gz). The boundary between the calyx and peduncle regions in the hemibrain was defined by innervation by PNs (or lack thereof) (personal communication, K. Shinomiya). The distance from this point to each PN-KC synapse along the KC’s neurite skeleton (i.e., not the Euclidean distance) was measured as described in [36].

## Code availability

Modeling and connectome analysis were carried out using custom code written in MATLAB, which is available at https://github.com/aclinlab/CompensatoryVariability.

## Acknowledgements

We thank Stuart Berg for ROI annotations in the connectome and Sofie Andersen, Caroni Fung, Celina Lubrino, Charles McMillan, and Edirimuni Rodrigo for help in identifying truncated KC skeletons. This work was supported by the European Research Council (639489 to ACL), the Biotechnology and Biological Sciences Research Council (BB/S016031/1 to ACL), and the Engineering and Physical Sciences Research Council (2131691 to NA, EV and ACL; EP/S009647/1, EP/S030964/1, EP/P006094/1 to EV).

## Author contributions

NA, Conceptualization, Software, Formal analysis, Investigation, Visualization, Methodology, Writing - original draft, Writing - review and editing. EV, Conceptualization, Formal analysis, Supervision, Funding acquisition, Methodology, Writing - review and editing. ACL, Conceptualization, Software, Formal analysis, Supervision, Funding acquisition, Investigation, Visualization, Methodology, Writing - original draft, Writing - review and editing.

## Supplementary Material

**Figure S1:**
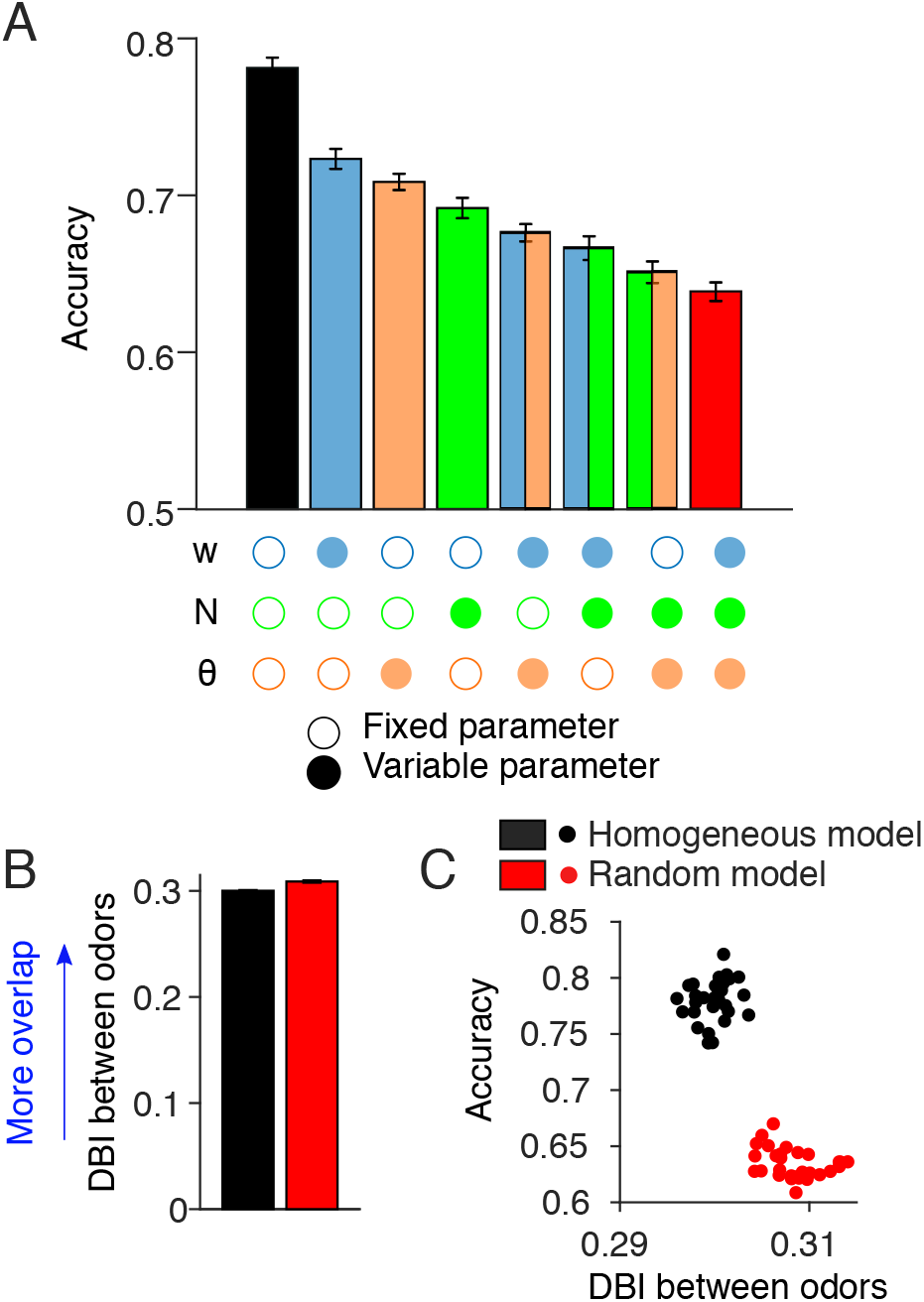
Similar analyses to Fig. 2 and 3 with original odor responses from [30]. **(A)** Inter-KC variability degrades the memory performance when using the 110 odorants from [30]. **(B-C)** The Davies-Bouldin index is higher in the random model.

**Figure S2:**
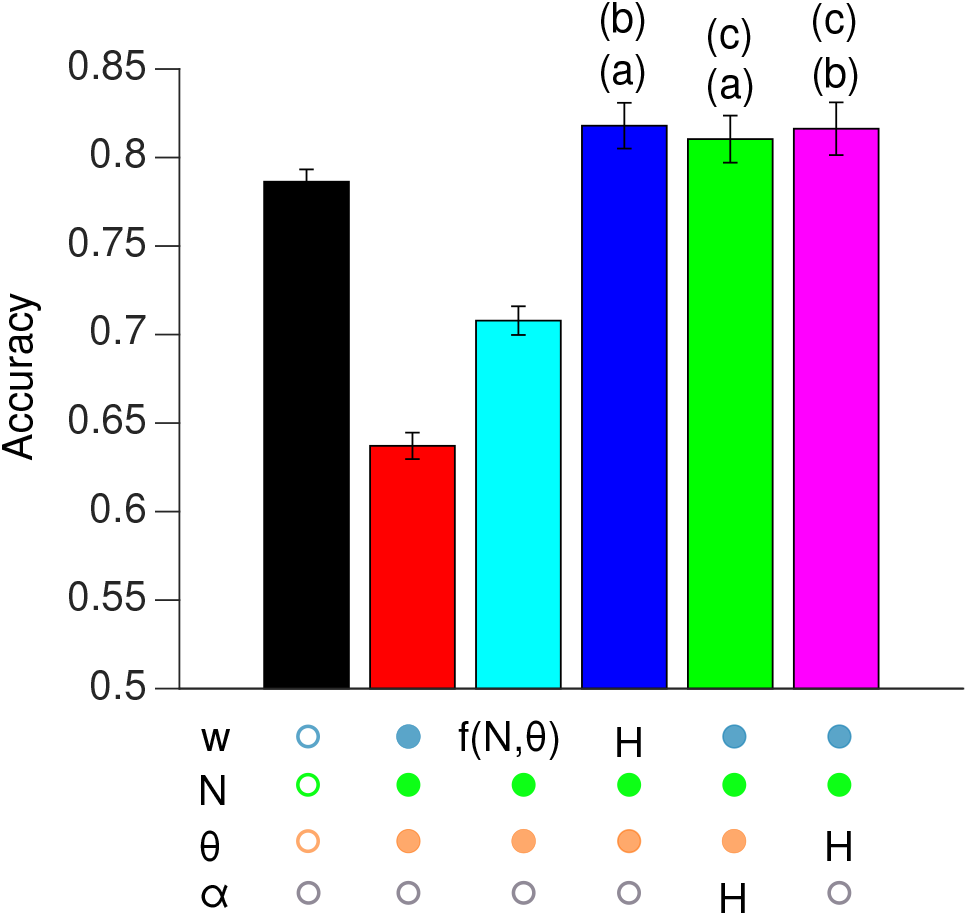
Similar analyses to Fig. 5 using the 110 odorants from [30]. The indeterminacy constant *c* from the softmax equation was set to 10. Bars that do not share the same letter annotation are significantly different, *p* < 0.05, Mann-Whitney or Wilcoxon test as in Fig. 5.

**Figure S3:**
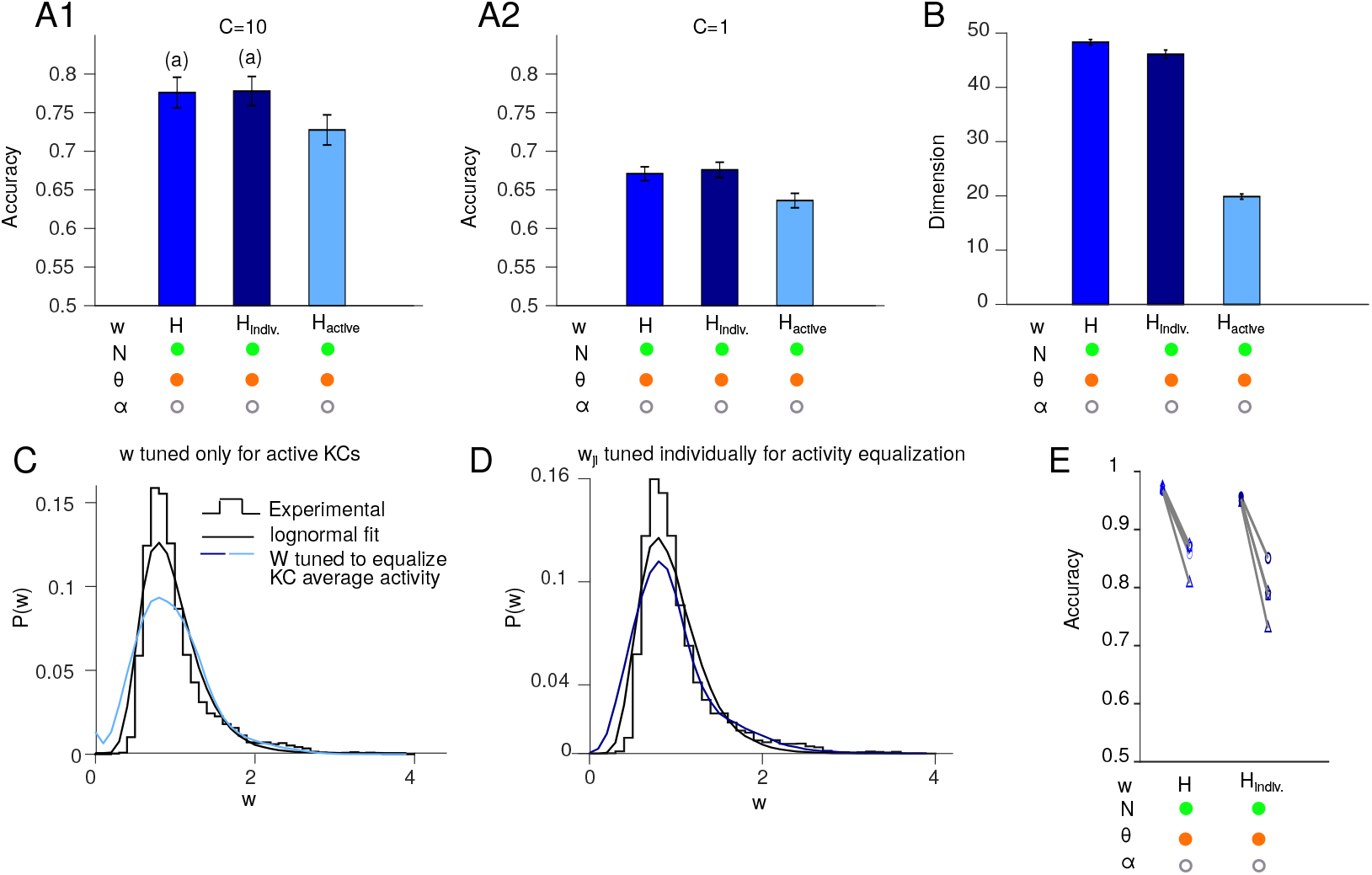
Alternative update rules for tuning KCs’ input excitatory weights. **(A)** Performance of different models at different indeterminacy constants (A1: *c* = 10; A2: *c* = 1): blue, the method in the main figures, Eq. (49), where a given KC’s input weights are all adjusted equally (‘H’); dark blue, Eq. (48), where a given KC’s input weights are adjusted individually according to the average activity of the PN (‘H_indiv_’); light blue, Eq. (46), where only non-silent KCs adjust their input weights (‘H_active_’). **(B)** Dimensionality of KC odor representations. The ‘H’ model has a significantly higher dimensionality than both the ‘H_indiv_’ and ‘H_active_’ models. *n* = 20 model instances with different random PN-KC connectivity. Error bars show two times the SEM, i.e., 95.4% confidence interval. Bars with the same letter annotations are not significantly different from each other; all other comparisons are significant *p* < 0.05, by Wilcoxon signed-rank test with Holm-Bonferroni correction for multiple comparisons. **(C-D)** Probability distribution of the tuned excitatory weights (compare to Fig. 5E). **(E)** The ‘H_indiv_’ model performs worse than the ‘H’ model in novel environments (see legend of Fig. 6).

**Figure S4:**
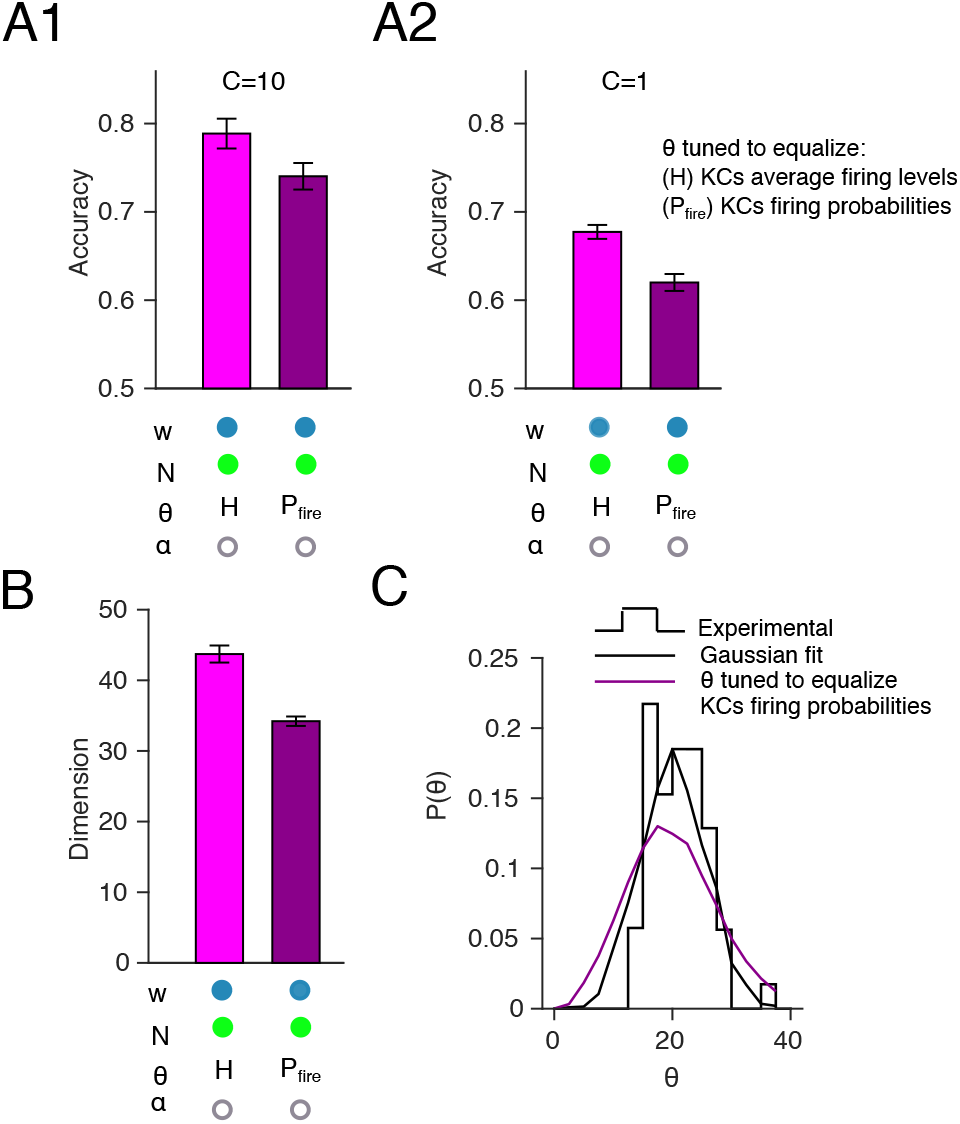
Equalizing KC average activity performs better than equalizing KC response probability. **(A)** Better performance when spiking thresholds are tuned to equalize KC average activity (magenta) rather than KC response probability (dark magenta), under both more (*c* = 10, A1) and less (*c* = 1, A2) deterministic decision-making. **(B)** Higher dimensionality of KC odor representations when equalizing KC average activity (magenta), compared to equalizing KC response probability (dark magenta). *n* = 20 model instances with different random PN-KC connectivity. Error bars show two times the SEM, i.e., 95.4% confidence interval. Magenta and dark magenta bars are significantly different, *p* < 0.05, by Wilcoxon signed-rank test. **(C)** Probability distribution of spiking thresholds (*θ*) after tuning them to equalize KCs’ response probabilities (compare to Fig. 5F).

**Figure S5:**
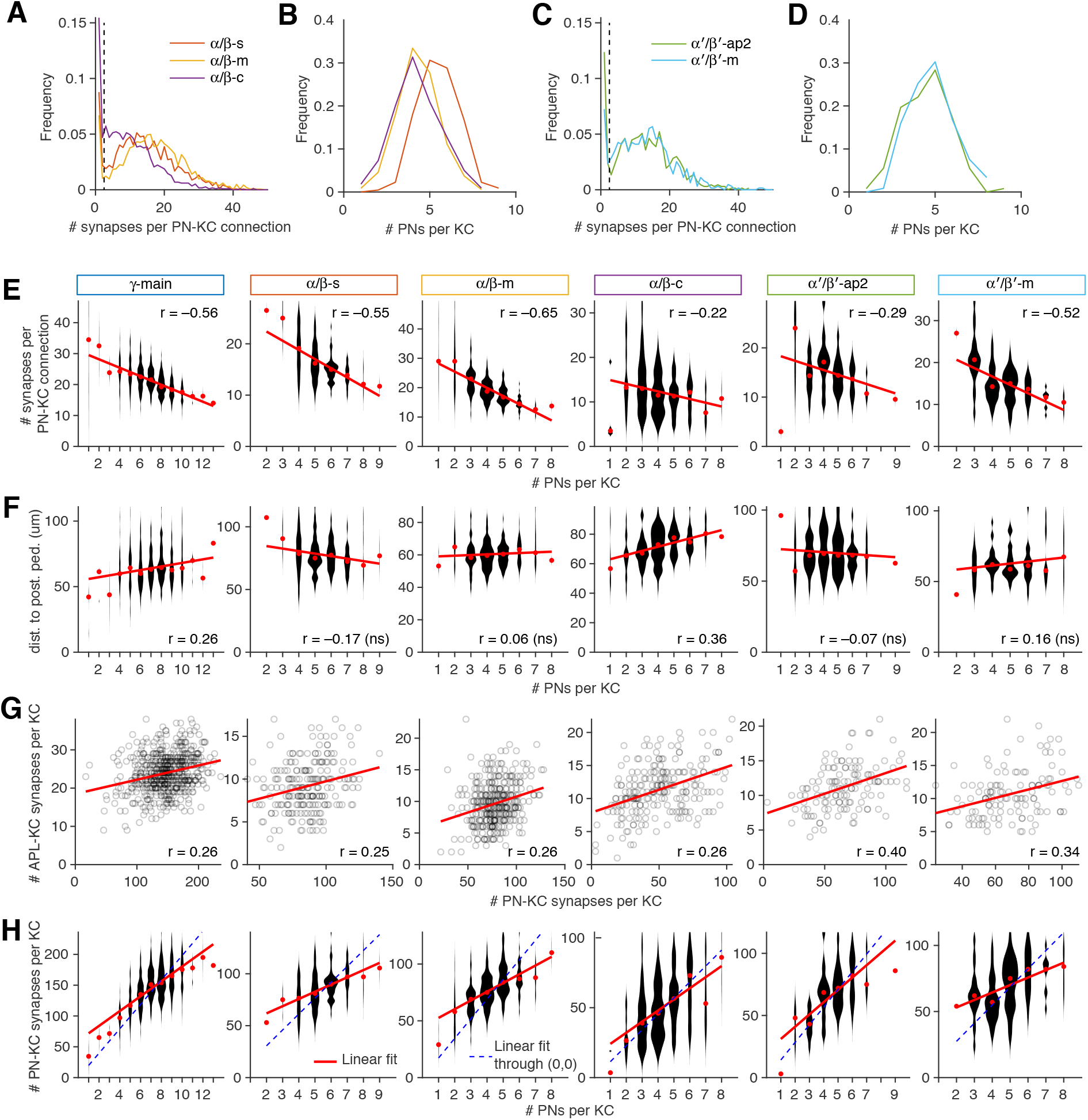
Connectome analysis on all KC subtypes (*γ*-main, *αβ*-s, -m and -c; *α′β′*-ap2 and -m). **(A-D)** Probability distributions of the number of synapses per PN-KC connection (A,C) and the number of input PNs per KC (B,D) in *αβ* and *α′β′* KCs separated out by subtype (compare to Fig. 7E,F). **(E)** Mean number of input synapses per PN-KC connection is inversely related to the number of input PNs per KC. **(F)** Mean distance of PN-KC synapses to the posterior boundary of the peduncle (presumed spike initiation zone) is directly related to the number of input PNs per KC in *γ* and *αβ*-c KCs. **(G)** The number of APL-KC synapses per KC is directly related to the total number of PN-KC synapses per KC. **(H)** The number of PN-KC synapses per KCs grows sublinearly with the number of PN inputs per KC. Red dots: medians. Red lines: linear fits. Blue dashed lines: linear fits through the origin (if every PN-KC connection had the same number of synapses). Note that the red dots follow a concave function relative to both linear fits.

